# Correlating basal gene expression across chemical sensitivity data to screen for novel synergistic interactors of HDAC inhibitors in pancreatic carcinoma

**DOI:** 10.1101/2022.10.10.511584

**Authors:** Nemanja Djokovic, Ana Djuric, Dusan Ruzic, Tatjana Srdic-Rajic, Katarina Nikolic

## Abstract

Pancreatic ductal adenocarcinoma (PDAC) is considered as one of the most aggressive and lethal malignancies. Development of chemoresistance in PDAC is one of the key contributors for the poor survival outcomes of PDAC patients and the major reason for urgent development of novel pharmacological approaches for effective treatment of PDAC. Systematically tailored combination therapy holds the promise for advancing the treatment of PDAC, but number of possible combinations considering all approved drugs and drug candidates is too large to be explored empirically. In respect to the many epigenetic alterations in PDAC, epigenetic drugs including histone deacetylase inhibitors (HDACi) could be seen as a game changer but available data indicates their efficacy only in combined therapy settings. In this work, we explored possibility of using drug-sensitivity data together with basal gene expression data on pancreatic cell lines to predict the combinatorial options available for HDACi and developed bioinformatics screening protocol for predictions of synergistic drug combinations in PDAC. Our results identified sphingolipid signaling pathway with associated downstream effectors as a promising novel target for future development of multi-target therapeutics or combined therapy with HDACi. Through the experimental validation, we have characterized novel synergism between HDACi and a Rho-associated protein kinase (ROCK) inhibitor RKI-1447, and HDACi and a sphingosine 1-phosphate (S1P) receptor agonist fingolimod.

## Introduction

Pancreatic ductal adenocarcinoma (PDAC) is considered as one of the most aggressive and lethal malignancies. Early onset dedifferentiation and metastasis significantly humper early diagnosis and treatment options which contributes to the devastatingly poor prognosis of the PDAC.^1–3^ Although, the surgical resection followed by adjuvant chemotherapy offers some prospects of long-term survival, over 80% of PDAC patients are diagnosed usually in terminal stages with distant metastasis have already occurred leaving the chemotherapy as the most frequently applied treatment option.^4–6^ Additionally, the most of the patients who have received surgery suffer from recurrence within a year, which makes the chemotherapy the mainstay of PDAC therapy.^6^ Considering the very modest improvement in 5-years survival rate since 1970s from 3% to 11%, there is an urgent need to develop non-surgical therapeutic approaches for effective treatment of pancreatic cancer.^7,8^

Gemcitabine, with or without additional chemotherapeutics, is considered as a first line option in chemotherapy of PDAC. Being one of the most chemoresistant cancers, efficacy of currently available chemotherapeutics for PDAC is seriously compromised by promptly developing chemoresistence.^2,6^ Dense stromal environment, many genetic and epigenetic alterations are evidenced to contribute to the complex mechanisms of the resistance. Inspired by many epigenetic alterations in PDAC, preclinical studies investigating interference with epigenetic mechanisms underlying PDAC by epigenetic therapeutics resulted in promising findings. However, increasing amount of evidence suggests that epigenetic therapeutics offers measurable benefits in PDAC only in combined therapy settings.^5,8^ Histone deacetylase inhibitors (HDACi) represent some of the most promising epigenetic therapeutics for treatment of PDAC.^8^ Currently, there are four HDACi approved by US Food and Drug Administration (FDA) while many candidate drugs are currently in clinical trials. Despite many promising preclinical results of HDACi used as single agents, clinical trials investigating efficacy of HDACi in solid tumors gave rather disappointing results. Therefore, a major approach for future developments in HDACi-based therapy is oriented towards application of these potent anticancer agents in combination with other chemotherapeutics.^8,9^

Despite significant breakthroughs in the cancer research during the last decades, high mortality remains major issue which highlights the urgent need for novel therapeutic approaches to the PDAC. Combination therapy holds the promise for advancing treatment of PDAC, but number of possible combinations considering all approved drugs and drug candidates in clinical trials is too large to be explored empirically. In an emerging era of personalized medicine, bioinformatics approaches could offer more rational way to select the optimal combination of drugs for specific sub-population of patients. In this work, we explored possibility of using drug-sensitivity data together with basal gene expression data on pancreatic cell lines to predict the combinatorial options available for HDACi. Our results identified sphingolipid signaling pathway with associated downstream effectors as a promising target for combined therapy with HDACi or future development of multi-target therapeutics. Through the process of experimental validation of the results, we have characterized novel synergism between HDACi and Rho-associated protein kinase (ROCK) inhibitors.

## Results and Discussion

Pharmaco-transcriptomics relationships between basal gene expression and chemical sensitivity of pancreatic carcinoma cell lines (CCLs) were investigated using systematic correlation analysis of large publically available data sets. Overall, Genomics of Drug Sensitivity in Cancer (GDSC)^10^ and Cancer Therapeutics Response Portal (CTRP)^11^ chemical sensitivity data sets for 864 small molecules across up to 38 pancreatic CCL were correlated to the basal gene expression of 19221 genes obtained from the Cancer Cell Line Encyclopedia (CCLE)^12^. Pearson correlation coefficients have been calculated between area under the curve (AUC) values and basal expression data (as log2RSEM) for each transcript. Matrix of Pearson correlation coefficients was further normalized using Fisher’s *z*-transformation to adjust for variations in number of tested CCLs for each of analyzed molecules. Generally speaking, transcripts negatively correlated with chemical sensitivity data could be attributed to the sensitivity of the pancreatic CCLs towards corresponding small molecule, while the positively correlated transcripts could be attributed to the resistance of the CCLs. In alignment to this, transcripts corresponding to the Tukey’s outlies of each small molecule were further extracted and stored as a transcriptomic “fingerprints” of drug sensitivity or resistance and used for further predictions of possible novel small molecule combinations. In this work we have hypothesized that level of overlap between these fingerprints could be an indicator of synergism between small molecules and could indicates novel pharmacological targets involved in synergy. The general approach of the study is presented on the Figure 1.

**Figure 1.**
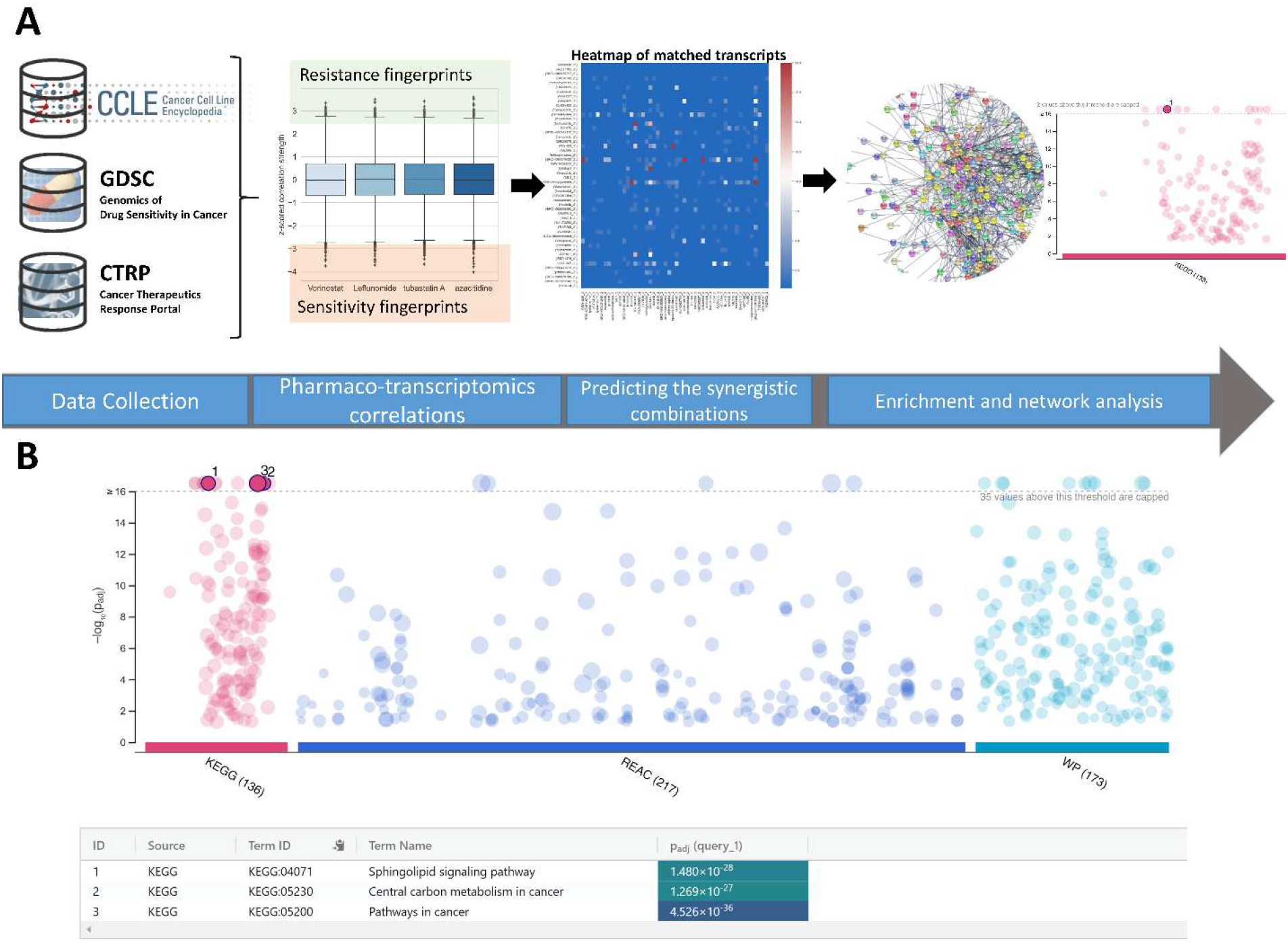
A – The preview of the approach used to identify synergistic interactors of HDACi and results of pathway enrichment analysis. B – Results of the enrichment analysis indicated sphingolipid signaling pathway as novel target for synergistic treatment of PDAC.

To further explore this hypothesis and identify novel potential synergistic counterparts to the HDACi, counting of the transcripts’ overlaps was performed on the pool of correlation data where only transcripts with significant correlation were considered. Namely, if the transcripts with significant positive correlation of first small molecules response had been overlapping with transcripts related to negative correlations of second small molecule response, these associations were counted and stored. Scores for all possible combinations of HDACi and the rest of small molecules had been calculated and probability of each of combination was evaluated as a count of all possible overlaps between significantly correlated transcripts (named “Final_Syn_Score” in the Supplementary Data File 1). Final results included 29233 possible combinations between HDACi and small molecule interactors (see Supplementary Data File 1).

Using the proposed protocol, several known synergistic interactors of HDACi have been identified which added up the validity of the approach. For example, the synergistic interaction between DNMT1 inhibitor and pan-HDACi was recently described on MiaPaCa2 cells.^13^ Sorafenib is one of the kinase inhibitors (BRAF; FLT3; KDR;RAF1) with preclinical data on synergism with HDACi and this combination is currently under clinical investigations for usage in PDAC^8,14^. Another positive examples include combination of HDAC inhibitors with PI3K inhibitor^15^, mTOR inhibitor^16^, BET (BRD4 in Table 1) inhibitors^17^, and combination with gemcitabine^9^ (Table 1). However, it should be noted that not all of the inhibitors of abovementioned pharmacological targets were highly scored in our analysis (for full table of predictions see Supplementary Data File 1) which could be attributed to the mostly unknown off-target effects of the small molecules.

**Table 1.**
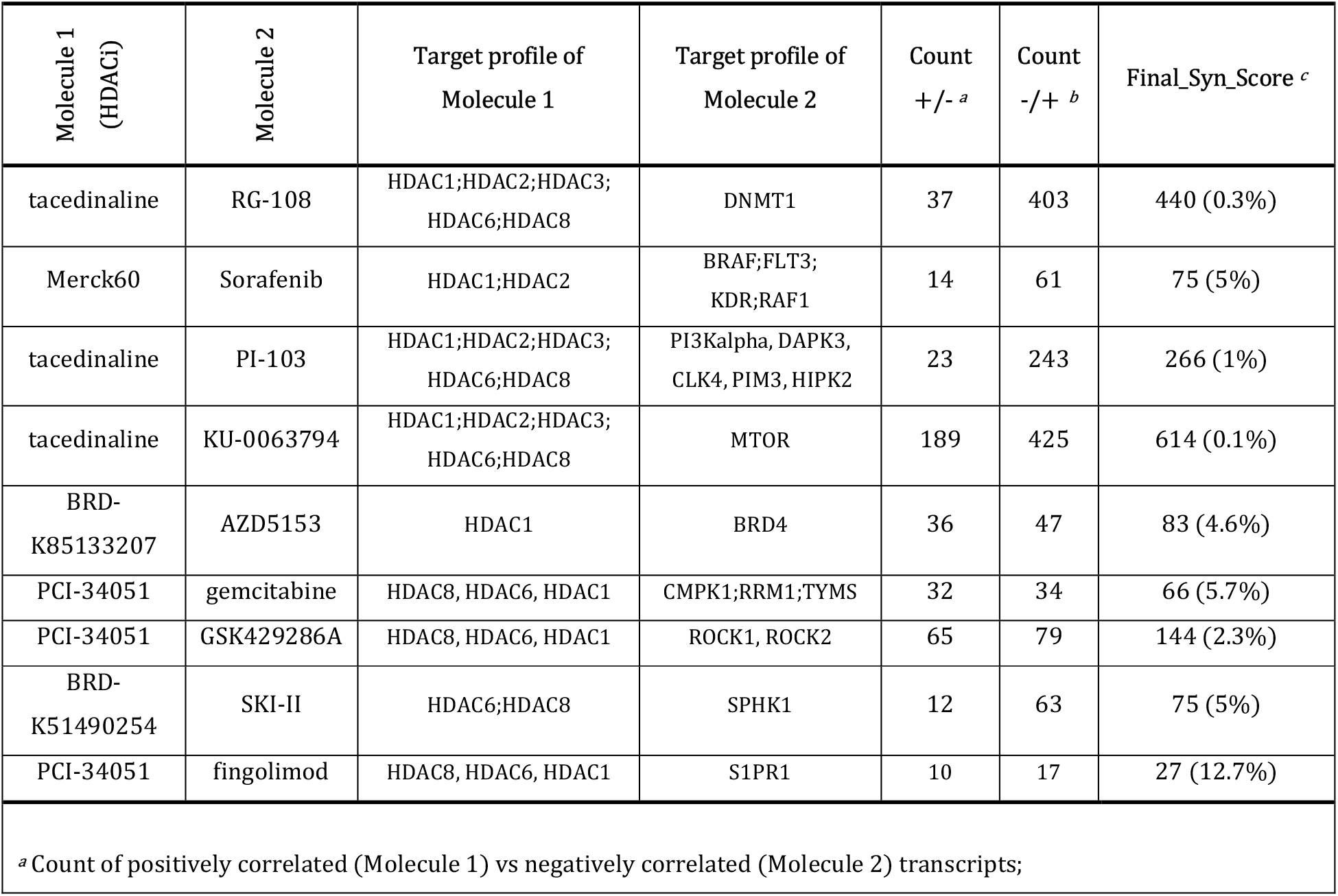
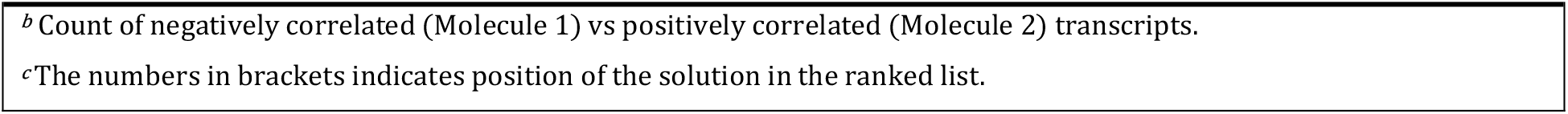
Results of HDACi synergy predictions with known literature data.

In order to identify possible signaling pathways which could be targeted for achieving synergistic effects with HDACi in PDAC, pathway enrichment analysis of annotated targets for predicted small molecule counterparts of HDACi was performed. Enrichment analysis revealed sphingolipid signaling pathway (KEGG)^18,19^ as one of the top-scored pathways involved in the predicted synergies across analyzed datasets of small molecules (Figure 1 and Supplementary Data File 2). Sphingolipid signaling pathway (as defined through analyzed KEGG and Wiki Pathway databases^20^) encompass several enzymes involved in sphingolipid metabolism (Sphingosine Kinase (SPHK), Ceramide Kinase (CERK), Ceramide Synthase (CERS), Alkaline Ceramidase (ACER), Acid Ceramidase (ASAH) etc.) as well as sphingosine-1-phosphate membrane receptors (S1PR1-5) and their downstream Rho/ROCK, PI3K/Akt and MAPK pathways. It is interesting to note that current literature data do not recognize any of the elements of sphingolipid signaling (except downstream pathways activated through many different pathways) as potential novel therapeutic targets for achieving synergism with HDACi which opens the great possibility for further research.

One of the central pillars of sphingolipid signaling represent so-called sphingolipid rheostat.^21^ Sphingolipid rheostat, as a concept within sphingolipid metabolism, could be seen as a dynamical equilibrium between amounts of ceramide which displays a pro-apoptotic role and sphingosine 1-phosphate (S1P) with a mitogenic and anti-apoptotic role. Sphingolipid rheostat, and particularly equilibrium between ceramide and S1P, have been just recently characterized as one of the critical regulator of pro- and anti-apoptotic singling in metabolically dynamic pancreatic cancers.^22,23^ Additionally, Speirs et al. identified SPHK1 as a key driver of the conserved S1P: Ceramide imbalance in pancreatic cancer subcultures and therefore one of the central controlling hubs of rheostat machinery.^22^ The most recent literature data suggested that sphingolipid signaling pathway could be an important novel therapeutic target to suppress proliferation across pancreatic tumors made up of heterogeneous cell populations, with recognition of potential of SPHK1 inhibitors as an approach to reverse healthy balance of pro- and anti-apoptotic signaling in pancreatic cancers.^22,24–26^ Additionally, predictions identified synergism between pan-HDAC inhibitor and SPHK1 inhibitor to be highly scored (in first 5% of the ordered list of predictions) (Table 1 and Supplementary Data File 1).

To further corroborate our results on potential synergism occurring between HDACi and sphingolipid signaling pathway in pancreatic carcinoma, expression analysis was performed on data obtained from pancreatic cancer cohort (The Cancer Genome Atlas (TCGA) database – number of samples 179) and healthy control (Genotype-Tissue Expression (GTEx) database – number of sample 171) (Figure 2). Besides overexpression of class I HDACs in pancreatic carcinoma patients, results indicated significant elevation of some of the key elements of sphingolipid signaling, including the enzymes involved in sphingolipid rheostat. Namely, expression of SPHK1, but not SPHK2, in patient tissue was significantly increased. Additionally, according to the differential expression analysis (Figure 2), expression of many components of the sphingolipid signaling pathway were significantly elevated in pancreatic cancer tissue, with most of them being upregulated. In summary, expression analysis suggested significant perturbations of sphingolipid signaling pathway in pancreatic carcinoma and together with overexpression of HDACs and synergy predictions further supports exploration for dual targeting of HDACs and sphingolipid signaling as novel approach in treatment of PDAC (Figure 2).

**Figure 2.**
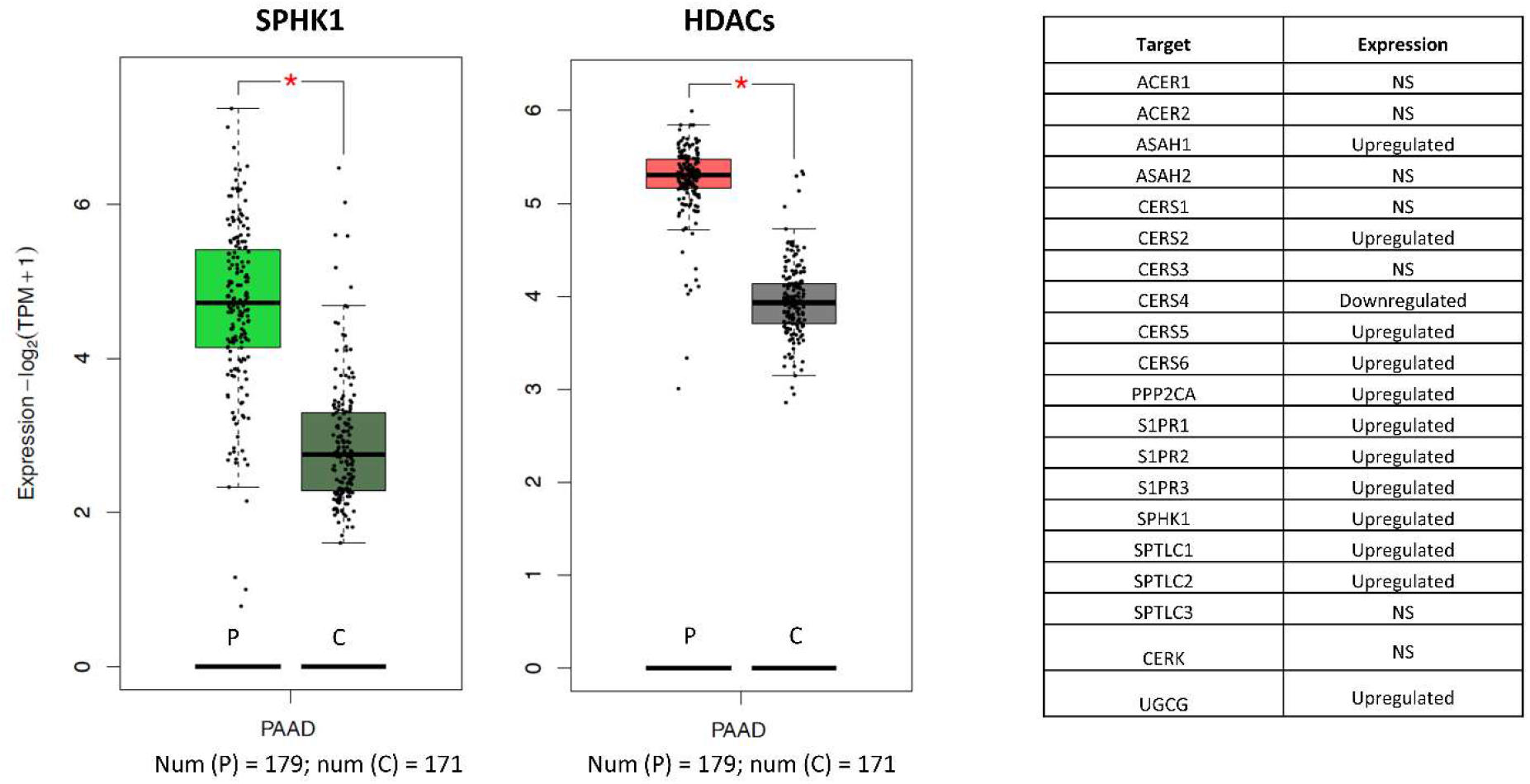
SPHK1 and HDACs (class I) are upregulated in pancreatic cancer. Left – Box plot of the expression for the central regulator of sphingolipid rheostat; Middle – Box plot of the expression of class I HDACs; Right – Table of summarized expressions for all component of sphingolipid signaling pathway identified in the study. Letter P indicates patient tissue, C indicates healthy control, while NS indicates non-significant change in expression between patient and normal tissue.

As a final step of our bioinformatics analysis, interactions network of all of the overlapped transcripts identified for synergisms between HDACi and elements of sphingolipid signaling pathway was constructed. Under assumption that these overlapped transcripts could be related to the mechanisms of synergy happening on the protein level, networks were created utilizing STRING database^27^ integrating all known and predicted associations between proteins, including both physical interactions as well as functional associations. Aiming to find underlying mechanism of synergy, networks were constructed using overlapped transcripts as nodes and STRING annotation on interactions as edges. Betweenness Centrality (BC) measurements were performed on created networks to identify the most important communication hubs inside networks, e.g. the nodes with the largest number of intersections of the shortest communication paths between other nodes. As a node with the highest BC value emerged p53 tumor suppresor, indicating importance of p53 in the networks mediating the synergistic effects between HDACi and modulators of sphingolipid signaling.

Through the complex interactions with HDACs including the post-translational regulation of acetylation status, p53 was found to be an important regulator of HDACi-mediated cancer cell death^28^. Additionally, literature data indicates that pro-apoptotic ceramide accumulation is an important downstream mediator of the p53 response further corroborating involvement of p53 in a mechanism of potential synergy between HDAC inhibition and modulation of sphingolipid signaling ^29^. Another identified communication hubs with corresponding BC values are presented in the Table 2.

**Table 2.** Communication hubs identified in network of transcripts involved in predicted synergism between HDACi and sphingolipid signaling pathway.

| Gene ID | BC | Gene ID | BC |
| --- | --- | --- | --- |
| TP53 | 56527 | APOA1 | 8858 |
| VWF | 19076 | ETV1 | 8308 |
| CALR | 17531 | PEBP1 | 7894 |
| NTRK1 | 16901 | ITGA4 | 7747 |
| MMP9 | 16341 | GNGT2 | 7426 |
| CYCS | 13336 | SLC17A7 | 7102 |
| DNM1 | 11804 | TPT1 | 6997 |
| FOXP3 | 10884 | BUD13 | 6632 |
| H2AFX | 10415 | COPS5 | 6290 |
| RCC1 | 9091 | DDB2 | 6156 |

In order to experimentally evaluate predictive power of proposed approach, synergisms between novel HDAC inhibitors recently described by our group together with modulators of pharmacological targets implicated by bioinformatics analysis available in the laboratory at the time of the study have been analyzed. For this analysis we have used a set of recently developed HDAC inhibitors: Compound 6b (IC_50_ (HDAC1) = 4.73 μM, IC_50_ (HDAC3) = 1.86 μM, IC_50_ (HDAC6) = 0.19 μM, IC_50_ (HDAC8) = 2.44 μM); Compound 8b (IC_50_ (HDAC1) = 0.41 μM, IC_50_ (HDAC3) = 0.21 μM, IC_50_ (HDAC6) = 0.12 μM, IC_50_ (HDAC8) = 0.24 μM); Compound 9b (IC_50_ (HDAC1) = 1.47 μM, IC_50_ (HDAC3) = 3.47 μM, IC_50_ (HDAC6) = 0.03 μM, IC_50_ (HDAC8) = 0.71 μM)^30^.

One of the pairs listed among 2.3% of sorted solutions was the potential synergy between Rho-associated protein kinase (ROCK) 1/2 inhibitors and pan-HDACi (Table 1 and Supplementary Data File 1). Interestingly, ROCK inhibi-tion, besides its effects on invasion and tumor growth, was previously described as valuable “priming” strategy for chemotherapy of PDAC ^31,32^. ROCK is also identi-fied as one of the downstream effectors of the sphingolipid S1P signaling ^19,33^. In experimental analysis we have used ROCK1/2 inhibitor RKI-1447 (IC_50_ (ROCK1) = 14.5 nM, and IC_50_ (ROCK2) = 6.2 nM). Fingolimod - another compound predicted to be syn-ergistic counterpart of pan-HDACi, was available for experimental evaluation in the laboratory at the time of the study. Fingolimod, the FDA-approved agonist and func-tional antagonist of S1P receptor subtype 1 (S1PR1) ^34^, was found to be potent anti-cancer agent in some PDAC models which could be at least partially attributed to its moderate inhibitory effects of the class I of HDACs ^35,36^. Besides S1P receptors, an-other elements of sphingolipid signaling pathway were recognized as off-targets of fingolimod, including ceramide synthase 2 (CERS2), S1P lyase, and SPHK1. Altogether, fingolimod appears to be a remarkable chemical probe to tackle potential of synergism between HDACi and sphingolipid signaling modulation ^34^. However, it is important to note that fingolimod:HDACi pair was scored among first 13% of predictions indi-cating rather moderate, or less probable, synergy.

To assess the cytotoxic effects of the drug combinations, Chou-Talalay model was used, which requires drugs to be administered at a fixed dose ratio. MIA PaCa-2 and Panc-1 cells were treated with a combination of synthesized HDACi and RKI-1447, or HDACi and fingolimod in a 4 × 4 dose matrix and measured the resultant cell viabili-ties by the MTT assay (Supplementary Material, Supplementary Note S1). Isobolo-gram analysis of the drug combination treatment at low concentrations and concentrations up to the IC50 value revealed slight to moderate synergism for a combination of compound 6b and RKI-1447 as well as compound 9b and RKI-1447 on MIA PaCa-2 cells (Table 3). Synergistic effects of the combination treatment with compound 8b and RKI-1447 on cell viability were observed at the highest concentration (Table 3 and Supplementary Material, Supplementary Note S1). Considering the results on Panc-1 cell line, isobologram analysis of the drug combination treatment revealed slight synergism for the combination of compound 9b and RKI-1447 only at the lowest concentrations (Table 3 and Supplementary Material, Supplementary Note S1). Interestingly, synergism between HDACi and fingolimod was also observed dominantly in 6b and 9b (Table 4 and Supplementary Material, Supplementary Note S1). In alignment with predictions, these synergisms were moderate and observed mostly for the highest tested concentrations. Although the antagonisms were detected for some of the tested concentration pairs, these antagonisms are of less concern since the most of them were detected for lower fractions of the affected cell viability (third row of Tables 3 and 4 of each pair).

**Table 3.** Synergism analysis for combination of novel HDACi and RKI-1447.

|  | MIA PaCa-2 cell line |  |  |  | Panc-1 cell line |  |  |  |
| --- | --- | --- | --- | --- | --- | --- | --- | --- |
|  | Compound concentration <sup>a</sup> |  |  |  | Compound concentration <sup>a</sup> |  |  |  |
|  | I | II | III | IV | I | II | III | IV |
| HDACi (**6b**) | 48.60<br>±7.45 | 13.58<br>±2.33 | 4.80<br>±1.01 | 2.91<br>±0.87 | 23.61<br>±3.26 | 1.39<br>±0.55 | 0.47<br>±0.21 | 0.5<br>±0.2 |
| ROCKi (RKI-1447) | 64.18<br>±8.12 | 40.49<br>±5.55 | 37.49<br>±3.02 | 30.84<br>±7.33 | 62.76<br>±11.23 | 33.89<br>±8.92 | 22.02<br>±7.22 | 18.11<br>±5.90 |
| **6b** + RKI-1447 | 83.28<br>±10.34 | 55.48<br>±7.51 | 37.69<br>±5.38 | 38.31<br>±4.56 | 69.79<br>±10.45 | 48.51<br>±6.34 | 24.98<br>±3.13 | 8.92<br>±4.21 |
| Combination Index (CI) | 0.445 | 0.981 | 1.253 | 0.605 | 0.992 | 1.050 | 1.334 | 2.117 |
| Interaction **6b** + RKI-1447 <sup>b</sup> | +++ | + | -- | ++ | ± | ± | -- | --- |
| HDACi (**8b**) | 27.30<br>±4.44 | 1.83<br>±0.70 | 0.94<br>±0.45 | 0.57<br>±0.19 | 49.78<br>±6.77 | 38.21<br>±5.89 | 17.13<br>±2.11 | 0.1<br>±0.05 |
| ROCKi (RKI-1447) | 62.94<br>±10.11 | 42.93<br>±6.99 | 36.07<br>±7.51 | 35.46<br>±4.09 | 58.78<br>±5.55 | 30.77<br>±3.45 | 17.27<br>±2.28 | 7.41<br>±2.56 |
| **8b** + RKI-1447 | 73.02<br>±7.44 | 45.32<br>±5.14 | 31.14<br>±5.21 | 28.91<br>±2.34 | 63.94<br>±6.33 | 42.96<br>±5.79 | 21.99<br>±3.45 | 6.79<br>±3.06 |
| Combination Index (CI) | 0.333 | 1.564 | 2.469 | 1.505 | 3.133 | 1.433 | 1.266 | 1.474 |
| Interaction **8b** + RKI-1447 <sup>b</sup> | +++ | --- | --- | --- | --- | -- | -- | --- |
| HDACi (**9b**) | 44.65<br>±3.39 | 18.25<br>±2.22 | 1.71<br>±0.67 | 0.46<br>±0.20 | 45.18<br>±4.55 | 23.14<br>±7.42 | 7.56<br>±3.71 | 1.12<br>±0.89 |
| ROCKi (RKI-1447) | 61.64<br>±7.78 | 43.28<br>±5.38 | 35.02<br>±3.25 | 29.16<br>±2.16 | 64.82<br>±6.41 | 37.92<br>±5.34 | 19.76<br>±2.30 | 9.41<br>±1.06 |
| **9b** + RKI-1447 | 78.03<br>±7.18 | 64.37<br>±6.78 | 41.91<br>±5.98 | 33.32<br>±4.21 | 64.68<br>±5.01 | 51.26<br>±7.20 | 33.14<br>±6.10 | 18.01<br>±4.02 |
| Combination Index (CI) | 0.513 | 0.577 | 0.964 | 0.803 | 1.705 | 1.207 | 0.975 | 0.823 |
| Interaction **9b** + RKI-1447 <sup>b</sup> | +++ | +++ | ± | ++ | --- | -- | ± | + |

<sup>a</sup> Concentrations are presented in descending order (I is the highest, and IV the lowest tested concentration). Exact concentrations for each compound are presented in Supplementary Material, Supplementary Note S1.
<sup>b</sup> Legend: (+ + + + +) Very strong synergism; (+ + +) Synergism; (+ +) Moderate synergism; (+) Slight synergism; (±) Nearly additive; (-) Slight antagonism; (- -) Moderate antagonism; (- - -) Antagonism; (- - - -) Strong antagonism; (- - - - -) Very strong antagonism.

**Table 4.**
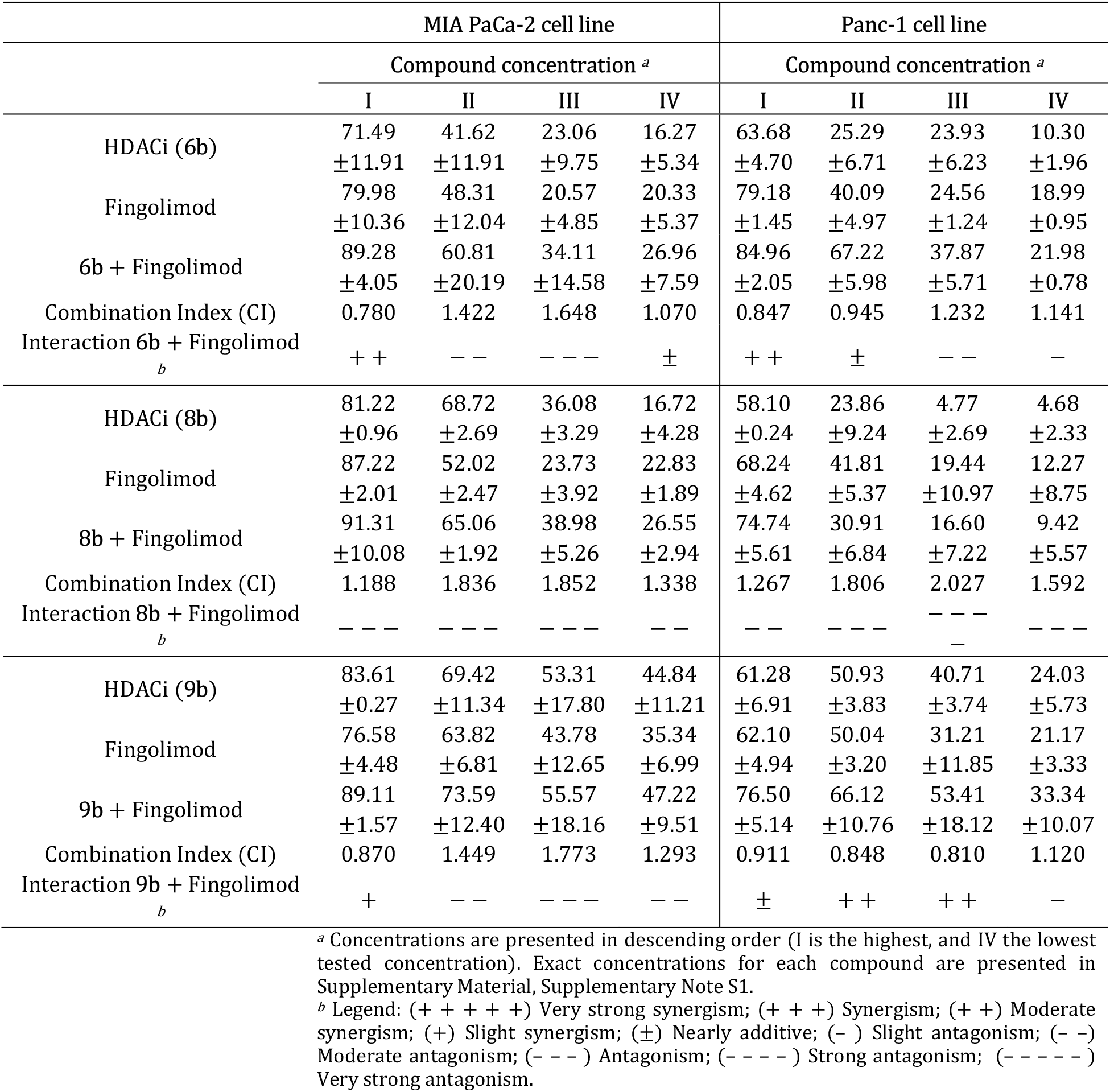
Synergism analysis for combination of novel HDACi and fingolimod.

Considering the fact that we were not tested exact predicted pairs of small-molecules, experimentally detected synergisms could indicate that these syner-gisms are driven by on-target rather than off-target effects of these drugs.

## Conclusion

Pancreatic ductal adenocarcinoma (PDAC) is considered as one of the most aggressive and lethal malignancies. Due to the high chemoresistance of PDAC as well as difficult early diagnosis which humpers surgical resection, there is an urgent need to develop novel pharmacological approaches for effective treatment of PDAC. In an emerging era of personalized medicine, systematically tailored combination therapy holds the promise for advancing treatment of PDAC, but number of possible combinations considering all approved drugs and drug candidates in clinical trials is too large to be explored empirically. In respect to the many epigenetic alterations in PDAC, epigenetic drugs including HDAC inhibitors could be seen as game changers. In this work, we explored possibility of using drug-sensitivity data together with basal gene expression data on pancreatic cell lines to predict the combinatorial options available for HDACi and developed bioinformatics screening protocol for predictions of synergy in PDAC. Our results identified sphingolipid signaling pathway with associated downstream effectors as a promising novel target for future development of multi-target therapeutics or combined therapy with HDACi. Through the process of experimental validation of the methodology, we have characterized novel synergism between HDACi and Rho-associated protein kinase (ROCK) inhibitors. The experimental validation of the protocol led to the discovery of the novel HDACi-ROCK inhibitor synergism as well as HDACi-fingolimod synergism. Further experimental evaluation of identified synergism between modulation of sphingolipid signaling and HDAC inhibition will be the major direction of our future work. All pre-dictions made through this study are freely available as a part of Supplementary Data File 1.

## Material and Methods

### Computational procedure

Pharmaco-transcriptomics relationships between basal gene expression and chemical sensitivity of pancreatic carcinoma cell lines (CCLs) were investigated using systematic correlation analysis of large publically available data sets: Genomics of Drug Sensitivity in Cancer (GDSC, https://www.cancerrxgene.org/)10 chemical sensitivity data sets, Cancer Therapeutics Response Portal (CTRP, https://portals.broadinstitute.org/ctrp.v2.1/)11 chemical sensitivity data set, Cancer Cell Line Encyclopedia (CCLE, https://depmap.org/portal/download/ 22Q2)^12^ basal gene expression data set. Pearson correlation coefficients have been calculated between area under the curve (AUC) values and basal expression data (as log2RSEM) for each transcript across all tested cell lines. Matrix of Pearson correlation coefficients was further normalized using Fisher’s *z*-transformation to adjust for variations in number of tested CCLs for each of analyzed molecules.. Transcripts corresponding to the Tukey’s outlies (1.5 interquartile range) of each small molecule were further extracted and stored as a transcriptomic “fingerprints” of drug resistance or sensitivity. Each of the transcripts from sensitivity or resistance signatures was further screened on the significance of positive or negative correlation coefficients across all analyzed small molecules. Only significant correlation were counted and each iteration was recorded as “Count +/-” or “Count -/+”. Total score, titled Final_Syn_Score, was calculated by summation across counts. List of annotated targets for all detected synergisms (Final_Syn_Score > 10 and at least one Tukey’s outlier transcript shared between small molecules) was further analyzed using the functional enrichment analysis from g:Profiler server (https://biit.cs.ut.ee/gprofiler)^37^. The Genotype-Tissue Expression (GTEx, https://www.gtexportal.org/home/) database and The Cancer Genome Atlas (TCGA, https://www.cancer.gov/tcga) were used to analyze the gene expression profiles of HDACs and elements of sphingolipid signaling across pancreatic carcinoma patient data. Data was accessed and analyzed through GEPIA web server (http://gepia.cancer-pku.cn/)^38^. Transcripts corresponding to the overlapped signatures identified across sphingolipid signaling pathway were extracted and analyzed through network analysis using the STRING^27^ data in Cytoscape 3.9.1.^39^ To identify major contributors to the communication across networks and mechanism of synergy, BC analysis was performed using CytoNCA tool.^40^

### Cell culture

MiaPaca-2 (ATCC CRL-1420), Panc-1 (ATCC CRL-1469) cells were cultured in DMEM medium (Sigma-Aldrich, St. Louis, MO, USA) supplemented with 10% fetal bovine serum (FBS) (Sigma-Aldrich, St. Louis, MO, USA), 100 μg/mL streptomycin and 100 UmL-1 penicillin (Sigma-Aldrich, St. Louis, MO, USA), and grown as a monolayer in humidified atmosphere of 95% air and 5% CO2 at 37 °C. in a 5% CO2 atmosphere at 370C and in humidified incubator.

### Drug interaction analysis

To determine the synergistic effects of the drug combinations, we performed an MTT viability assay and the combination index method described by Chou and Talalay.^41^ Cytotoxic activity of synthesized HDAC inhibitors and ROCK inhibitor was assessed on MiaPaCa-2 and Panc-1 cells using MTT assay.^42^ MiaPaCa-2 (4×103 cells/well) and PANC-1 (5×103 cells/well) were treated with synthesized compounds in five different concentrations (100, 50, 25, 12.5, and 6.25 μM), and each concentration is added in five replicates. After 72 h, 20 μL of MTT solution (3-(4, 5-dimethylthiazol-2-yl)-2, 5-dyphenyl tetrazolium bromide) (Sigma-Aldrich, St. Louis, USA) was added to each well. Samples were incubated for 4 h, followed by the addition of 100 μL of 10% SDS and incubated at 37 °C. Absorbance at 570 nm was measured the next day. Cell survival (%) was calculated as an absorbance (570 nm) ratio between treated and control cells multiplied by 100. IC_50_ was defined as the concentration of the agent that inhibited cell survival by 50% compared to the vehicle control.

To determine the synergistic effects of the drug combinations, we performed the combination index method described by Chou and Talalay^41^, using the CalcuSyn software (version 2.0 Biosoft, Cambridge, UK). The combination index (CI) was calculated to assess the benefits of combinational treatment in MiaPaCa-2 and PANC-1 cells. A CI equal to 1 indicates an additive effect, CI = 0.3–0.7 indicates synergism, CI = 0.1–0.3 indicates strong synergism, and CI < 0.1 indicates very strong synergism.

## Acknowledgments

N.D., D.R. and K.N. acknowledge the Ministry of Education, Science and Technological Development of the Republic of Serbia, Faculty of Pharmacy UB Contract No. 451-03-68/2022-14/200161 and COST actions CA18133 and CA18240.

## Author contributions

N.D. performed bioinformatics experiments and calculations. N.D., D.R. and K.N. conceived of the idea for the study, designed, directed and interpreted experiments. A.D. and T.S.R. performed experimental in vitro evaluation of synergisms. All of the authors contributed to the writing of the manuscript and the supplementary material.

