## Supplementary Material for "Correlating basal gene expression across chemical sensitivity data to screen for novel synergistic interactors of HDAC inhibitors in pancreatic carcinoma"

**Supplementary Note S1.** Detailed results of experimental evaluation of synergisms with concentration annotations.

**MiaPaCa2 cell line**

|  | Drug concentration | | | |
| --- | --- | --- | --- | --- |
| Drug | I | II | III | IV |
| HDACi (**6b**) | 48.60±7.45 | 13.58±2.33 | 4.80±1.01 | 2.91±0.87 |
| ROCKi (RKI-1447) | 64.18±8.12 | 40.49±5.55 | 37.49±3.02 | 30.84±7.33 |
| **6b** + RKI-1447 | 83.28±10.34 | 55.48±7.51 | 37.69±5.38 | 38.31±4.56 |
| Combination Index (CI) | 0.445 | 0.981 | 1.253 | 0.605 |
| Interaction **6b** + RKI-1447 | + + + | + | − − | + + |

Compound 6 (µM): I=50; II=25; III=12.5; IV=6.25

RKI (µM): I=30; II=15; III=7.5; IV=3.75

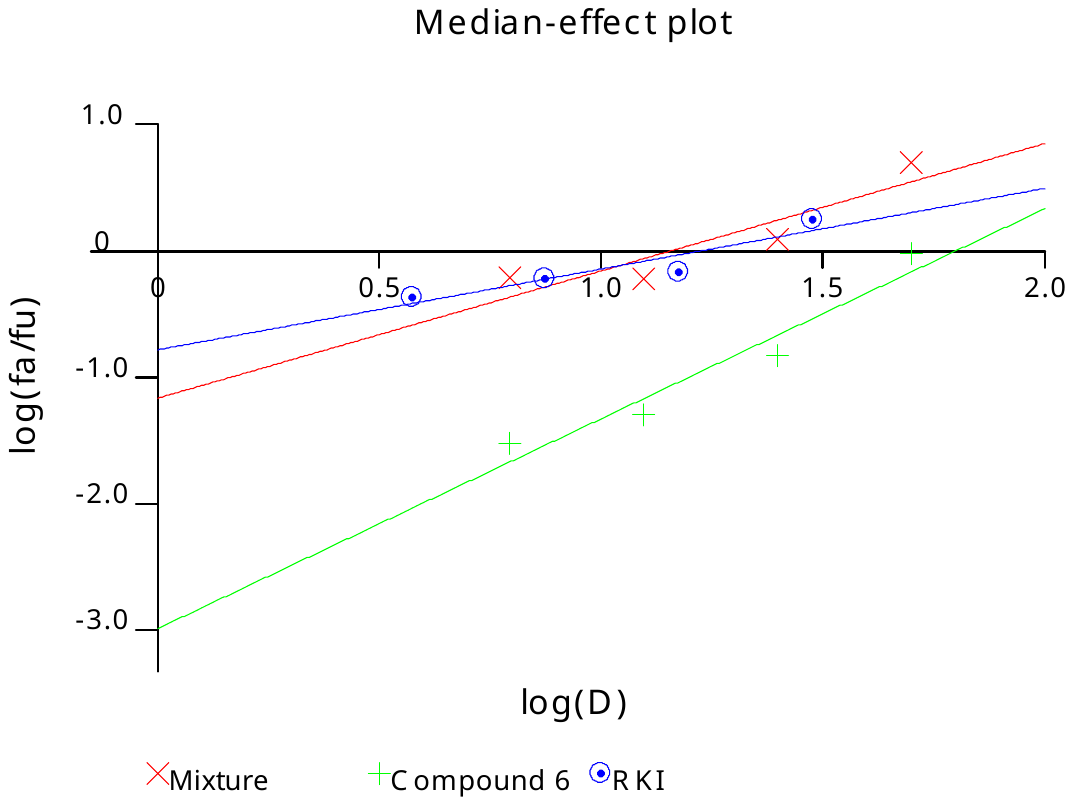

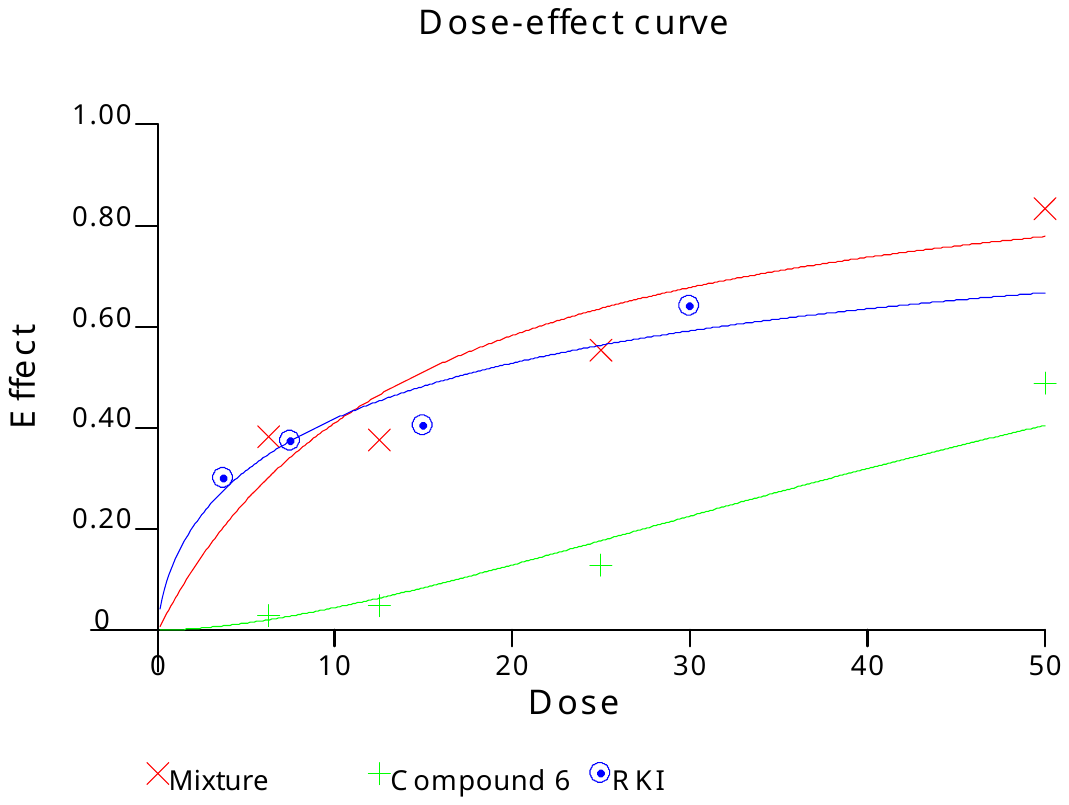

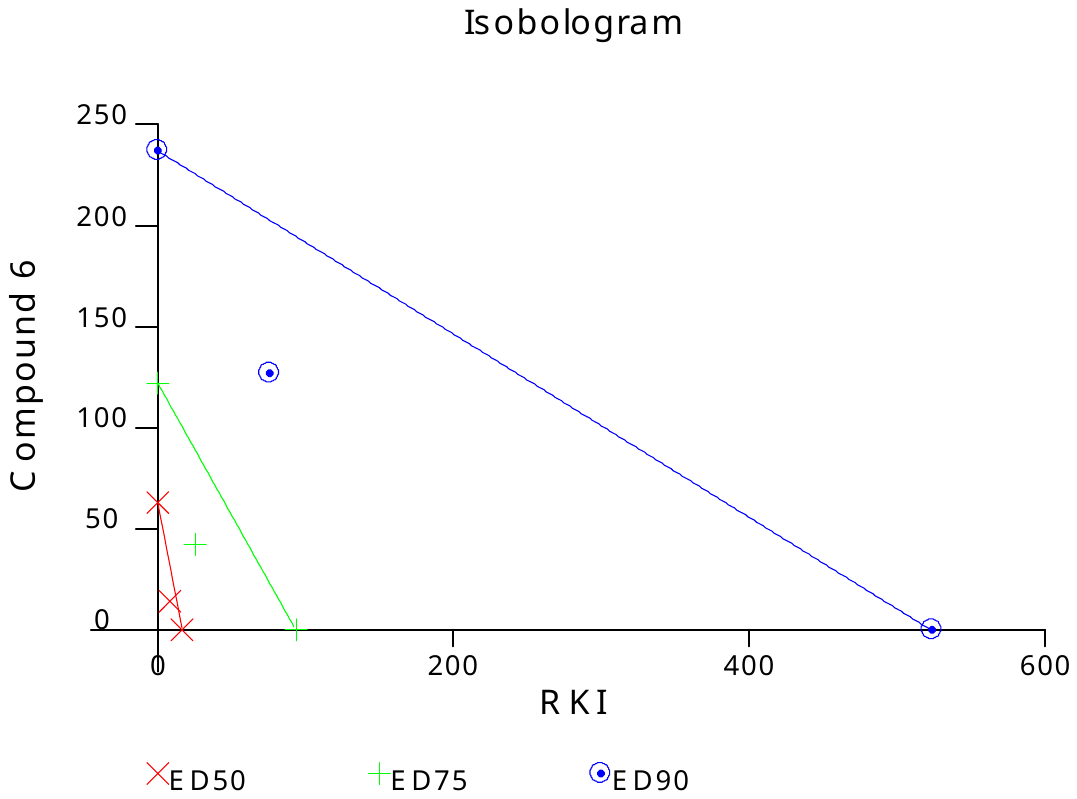

**MiaPaCa2 cell line**

|  | Drug concentration | | | |
| --- | --- | --- | --- | --- |
| Drug | I | II | III | IV |
| HDACi (**8b**) | 27.30±4.44 | 1.83±0.70 | 0.94±0.45 | 0.57±0.19 |
| ROCKi (RKI-1447) | 62.94±10.11 | 42.93±6.99 | 36.07±7.51 | 35.46±4.09 |
| **8b** + RKI-1447 | 73.02±7.44 | 45.32±5.14 | 31.14±5.21 | 28.91±2.34 |
| Combination Index (CI) | 0.333 | 1.564 | 2.469 | 1.505 |
| Interaction **8b** + RKI-1447 | + + + | − − − | − − − | − − − |

Compound 8 (µM): I=6; II=3; III=1.5; IV=0.75

RKI (µM): I=30; II=15; III=7.5; IV=3.75

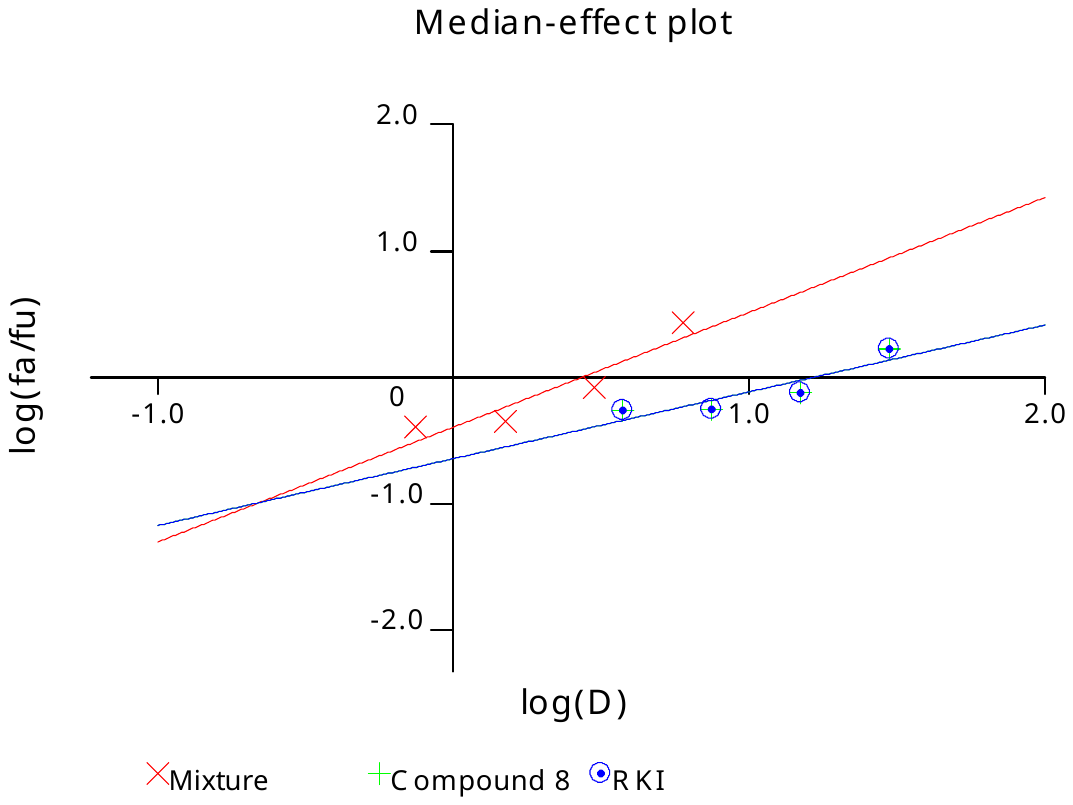

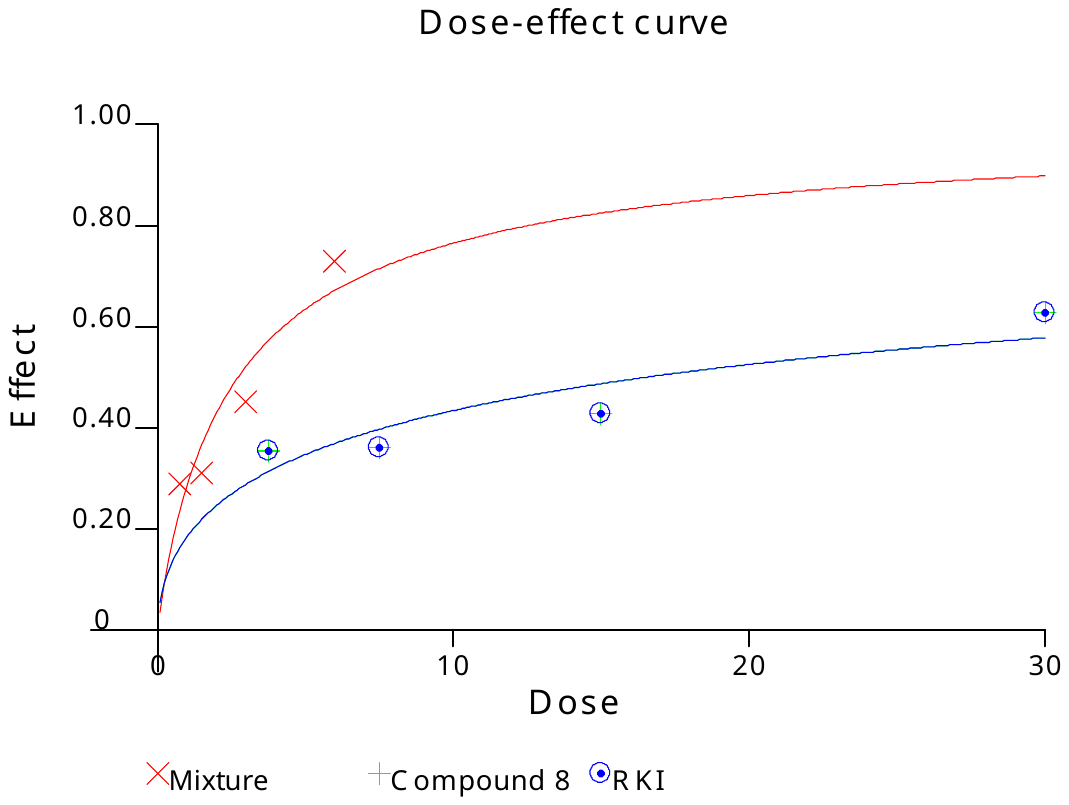

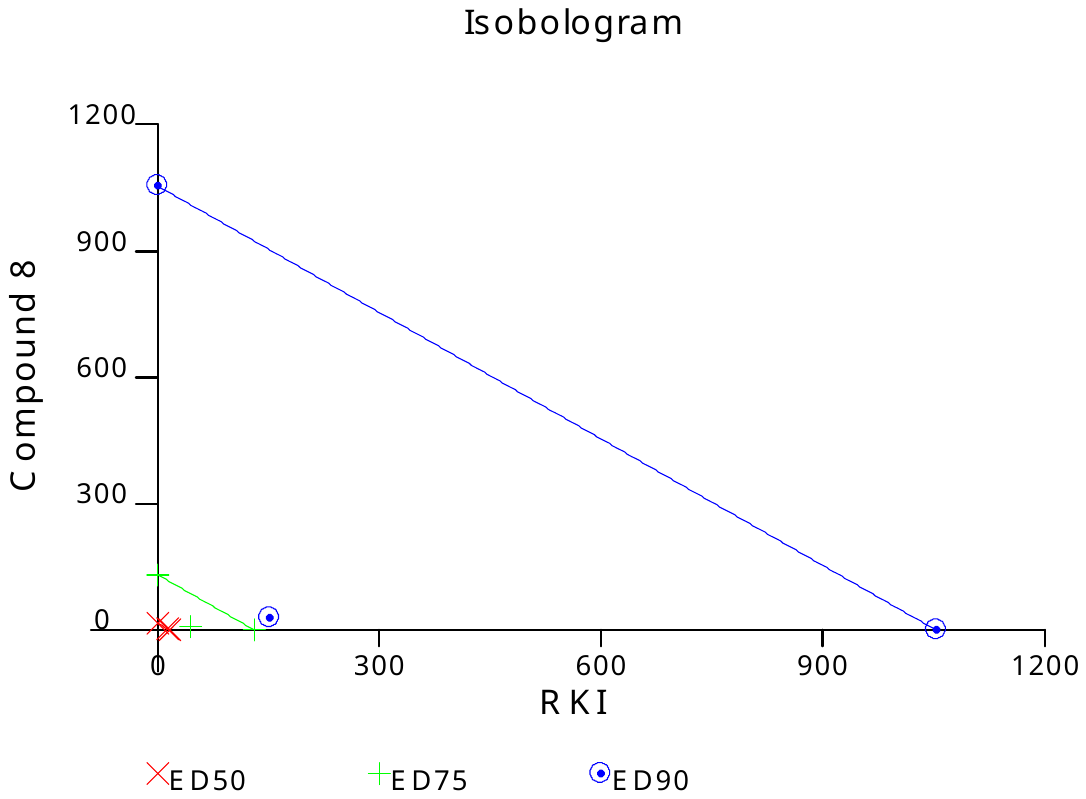

**MiaPaCa2 cell line**

|  | Drug concentration | | | |
| --- | --- | --- | --- | --- |
| Drug | I | II | III | IV |
| HDACi (**9b**) | 44.65±3.39 | 18.25±2.22 | 1.71±0.67 | 0.46±0.20 |
| ROCKi (RKI-1447) | 61.64±7.78 | 43.28±5.38 | 35.02±3.25 | 29.16±2.16 |
| **9b** + RKI-1447 | 78.03±7.18 | 64.37±6.78 | 41.91±5.98 | 33.32±4.21 |
| Combination Index (CI) | 0.513 | 0.577 | 0.964 | 0.803 |
| Interaction **9b** + RKI-1447 | + + + | + + + | ± | + + |

Compound 9 (µM): I=50; II=25; III=12.5; IV=6.25

RKI (µM): I=30; II=15; III=7.5; IV=3.75

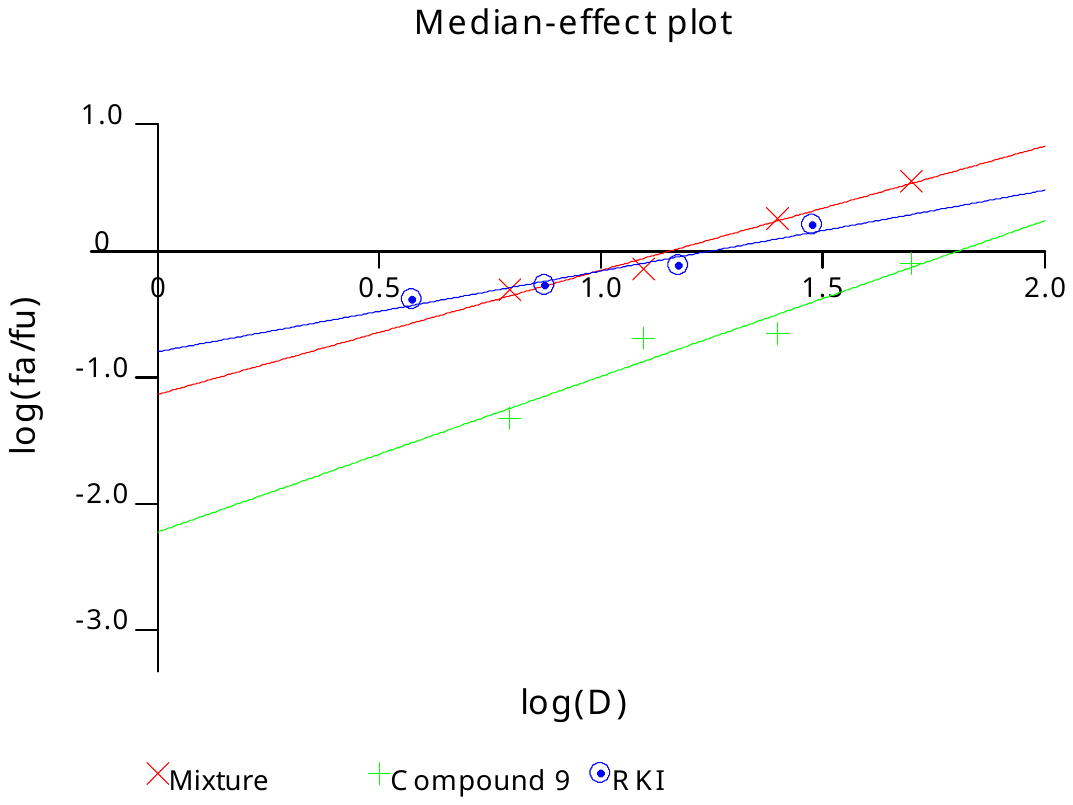

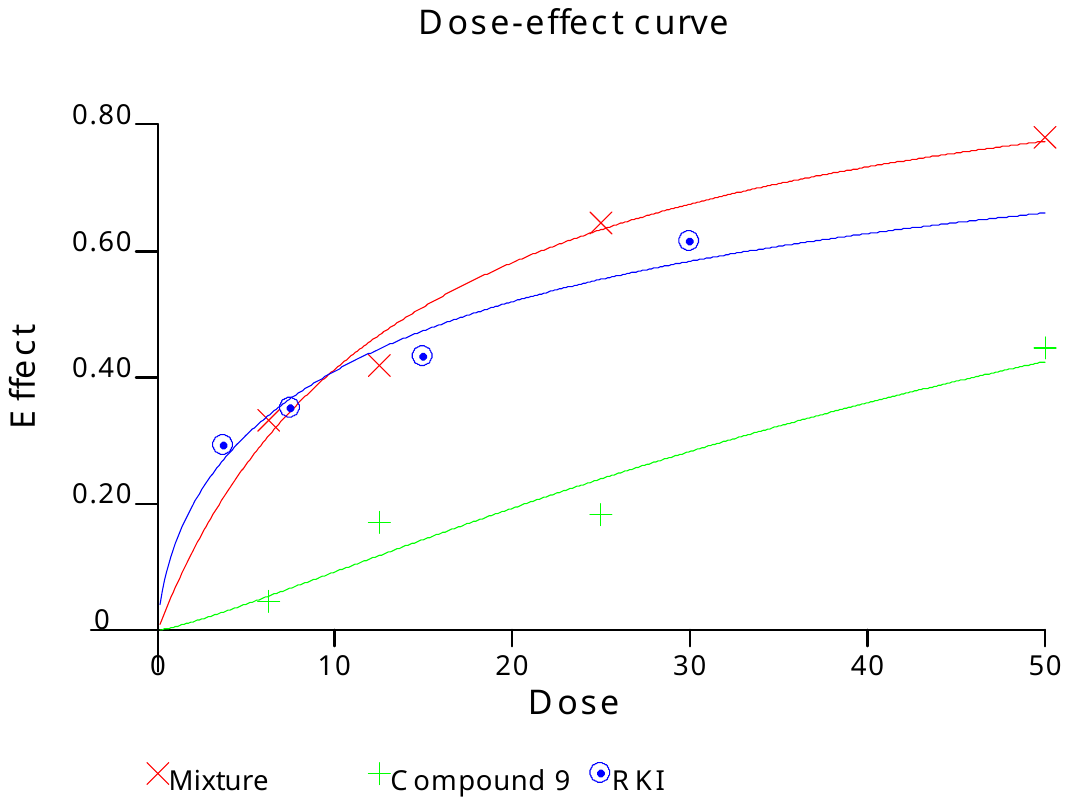

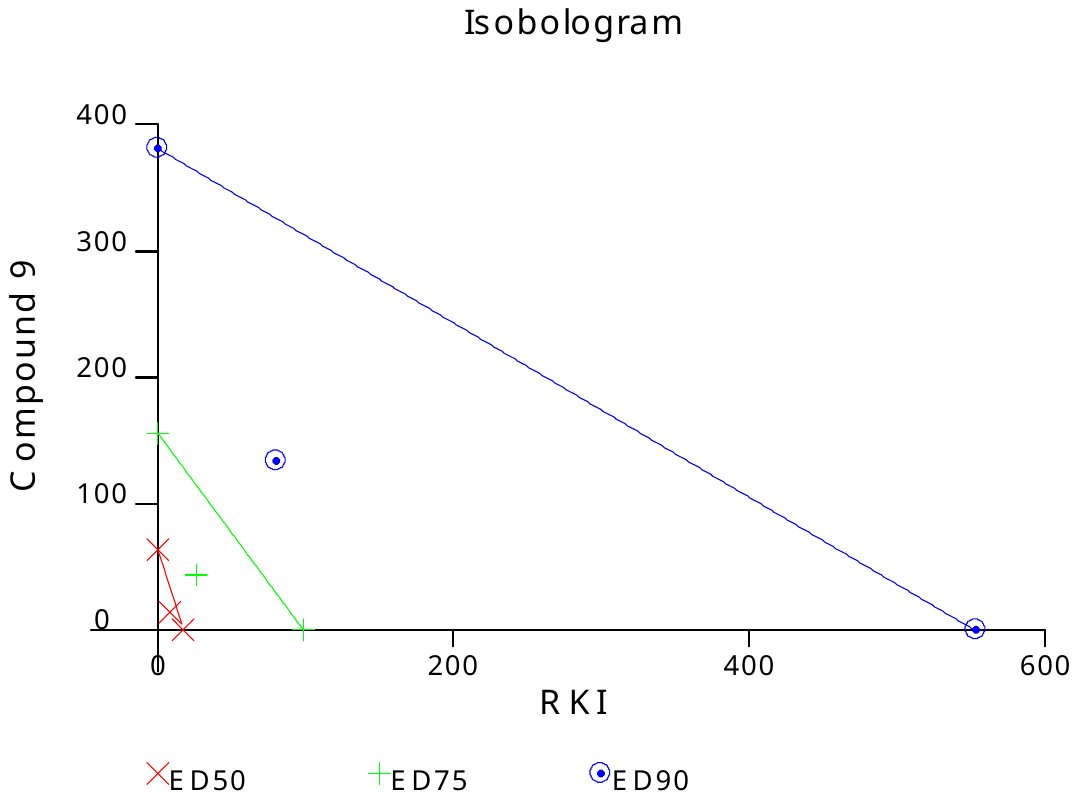

**Panc-1 cell line**

|  | Drug concentration | | | |
| --- | --- | --- | --- | --- |
| Drug | I | II | III | IV |
| HDACi (**6b**) | 23.61±3.26 | 1.39±0.55 | 0.47±0.21 | 0.5±0.2 |
| ROCKi (RKI-1447) | 62.76±11.23 | 33.89±8.92 | 22.02±7.22 | 18.11±5.90 |
| **6b** + RKI-1447 | 69.79±10.45 | 48.51±6.34 | 24.98±3.13 | 8.92±4.21 |
| Combination Index (CI) | 0.992 | 1.050 | 1.334 | 2.117 |
| Interaction **6b** + RKI-1447 | ± | ± | − − | − − − |

Compound 6 (µM): I=50; II=25; III=12.5; IV=6.25

RKI (µM): I=50; II=25; III=12.5; IV=6.25

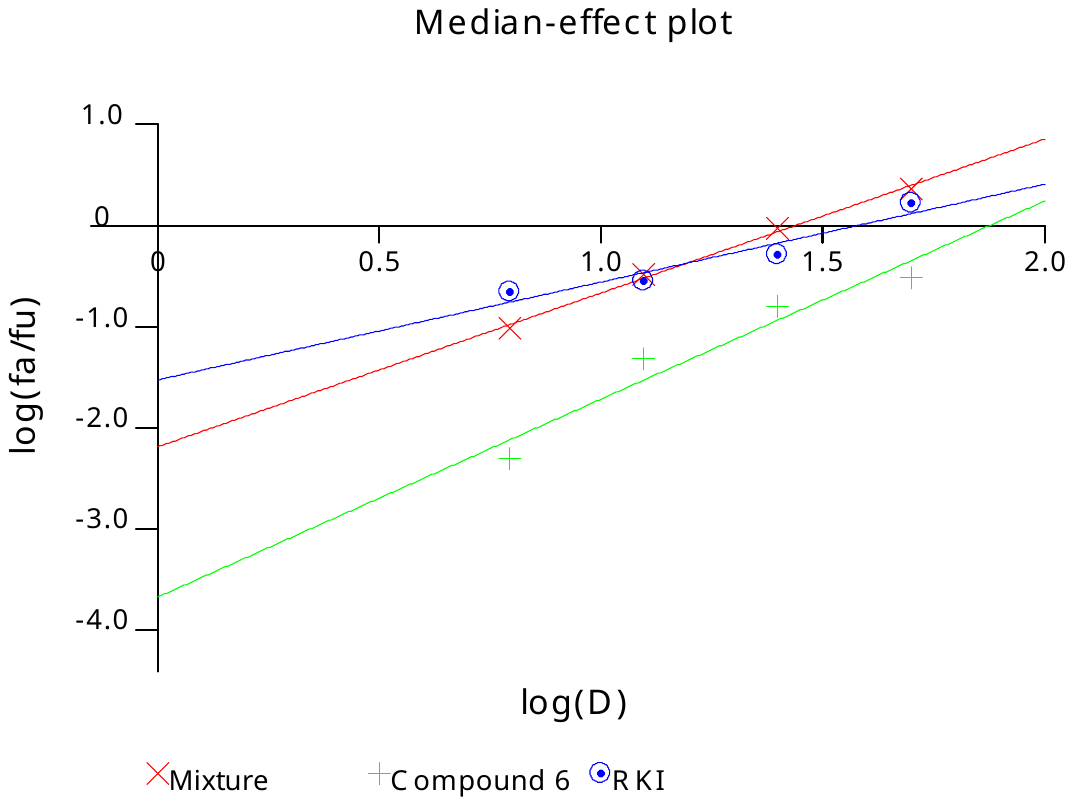

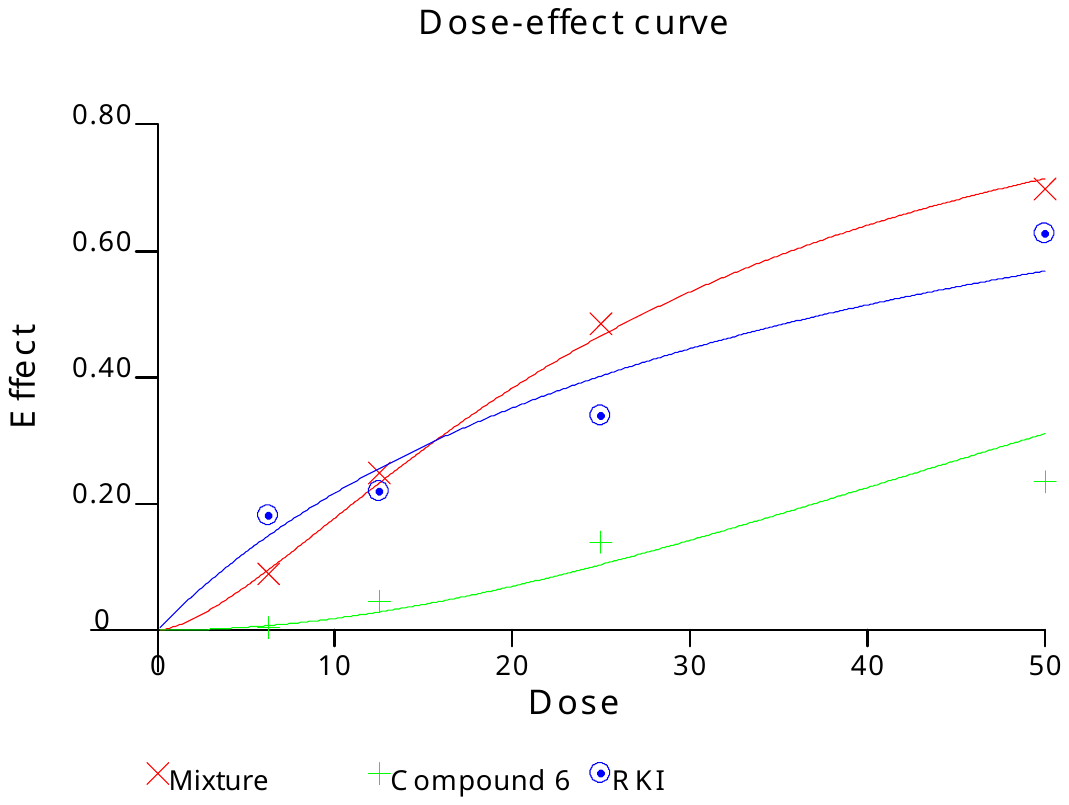

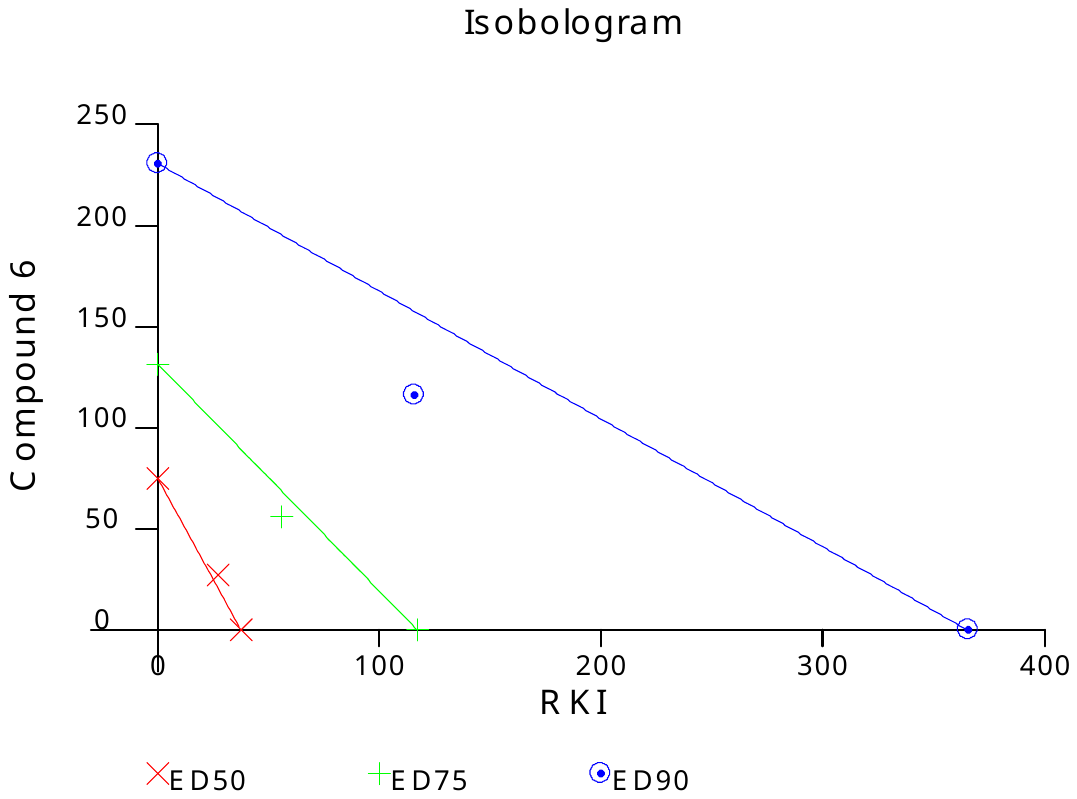

**Panc-1 cell line**

|  | Drug concentration | | | |
| --- | --- | --- | --- | --- |
| Drug | I | II | III | IV |
| HDACi (**8b**) | 49.78±6.77 | 38.21±5.89 | 17.13±2.11 | 0.1±0.05 |
| ROCKi (RKI-1447) | 58.78±5.55 | 30.77±3.45 | 17.27±2.28 | 7.41±2.56 |
| **8b** + RKI-1447 | 63.94±6.33 | 42.96±5.79 | 21.99±3.45 | 6.79±3.06 |
| Combination Index (CI) | 3.133 | 1.433 | 1.266 | 1.474 |
| Interaction **8b** + RKI-1447 | − − − | − − | − − | − − − |

Compound 8 (µM): I=20; II=10; III=5.0; IV=2.5

RKI (µM): I=50; II=25; III=12.5; IV=6.25

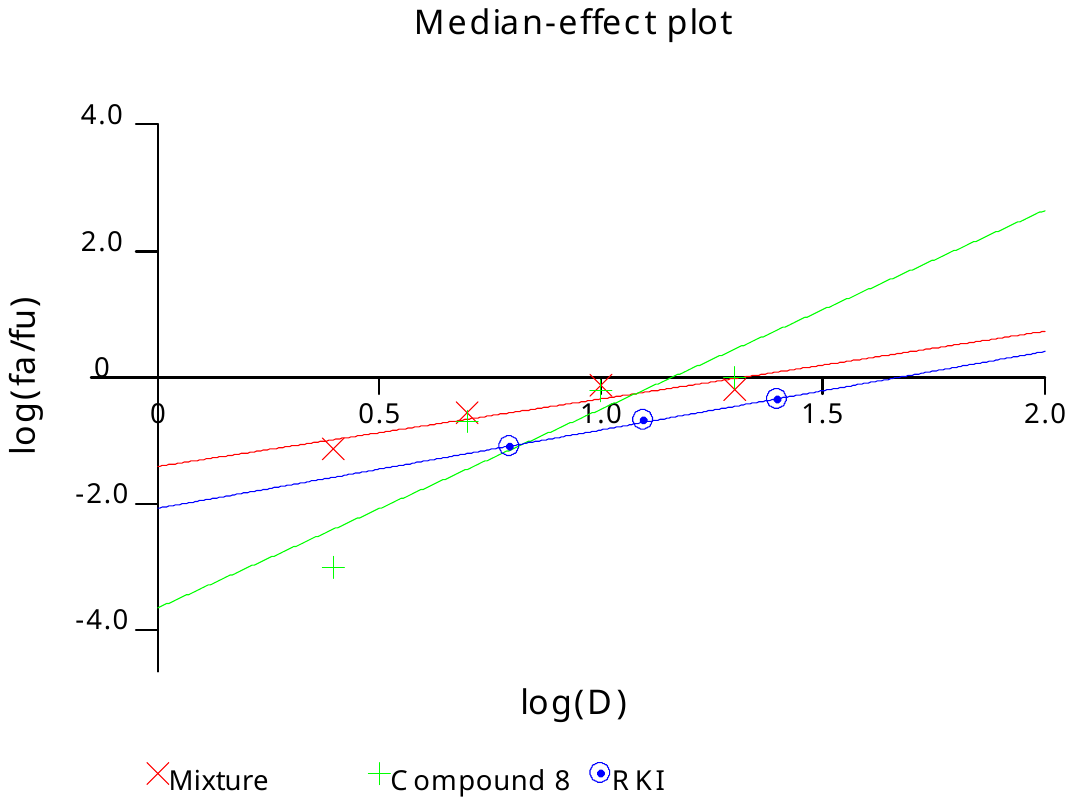

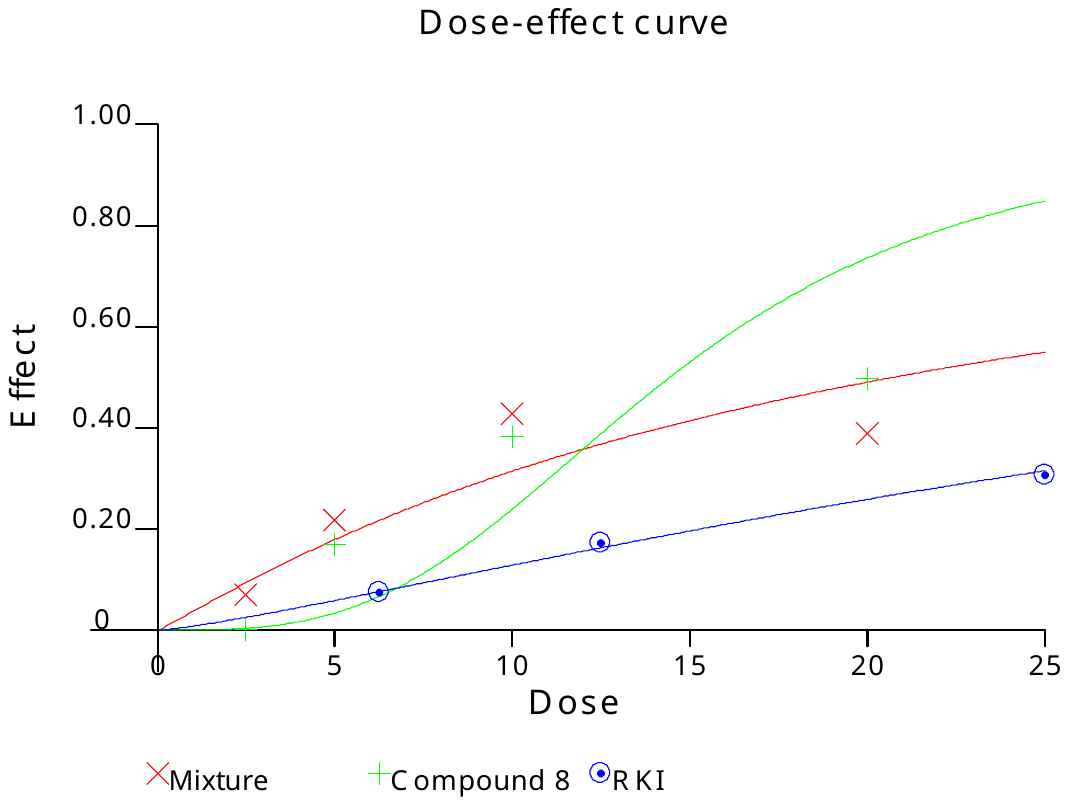

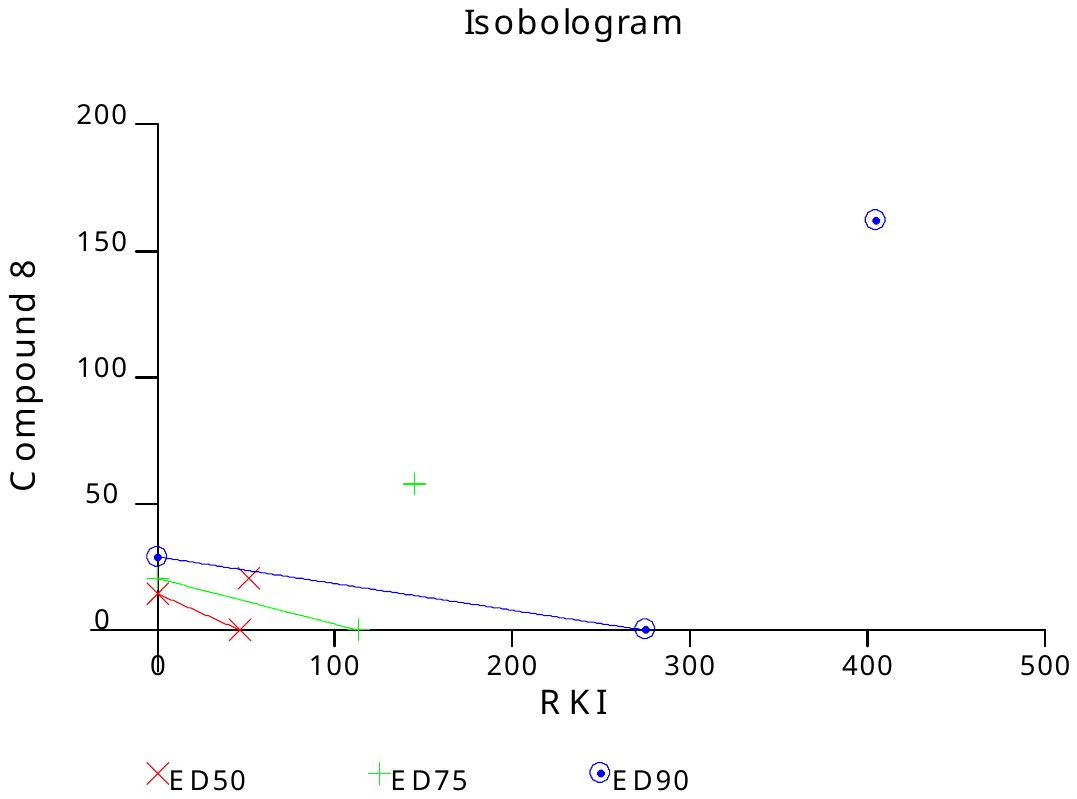

**Panc-1 cell line**

|  | Drug concentration | | | |
| --- | --- | --- | --- | --- |
| Drug | I | II | III | IV |
| HDACi (**9b**) | 45.18±4.55 | 23.14±7.42 | 7.56±3.71 | 1.12±0.89 |
| ROCKi (RKI-1447) | 64.82±6.41 | 37.92±5.34 | 19.76±2.30 | 9.41±1.06 |
| **9b** + RKI-1447 | 64.68±5.01 | 51.26±7.20 | 33.14±6.10 | 18.01±4.02 |
| Combination Index (CI) | 1.705 | 1.207 | 0.975 | 0.823 |
| Interaction **9b** + RKI-1447 | − − − | − − | ± | + |

Compound 9 (µM): I=50; II=25; III=12.5; IV=6.25

RKI (µM): I=50; II=25; III=12.5; IV=6.25

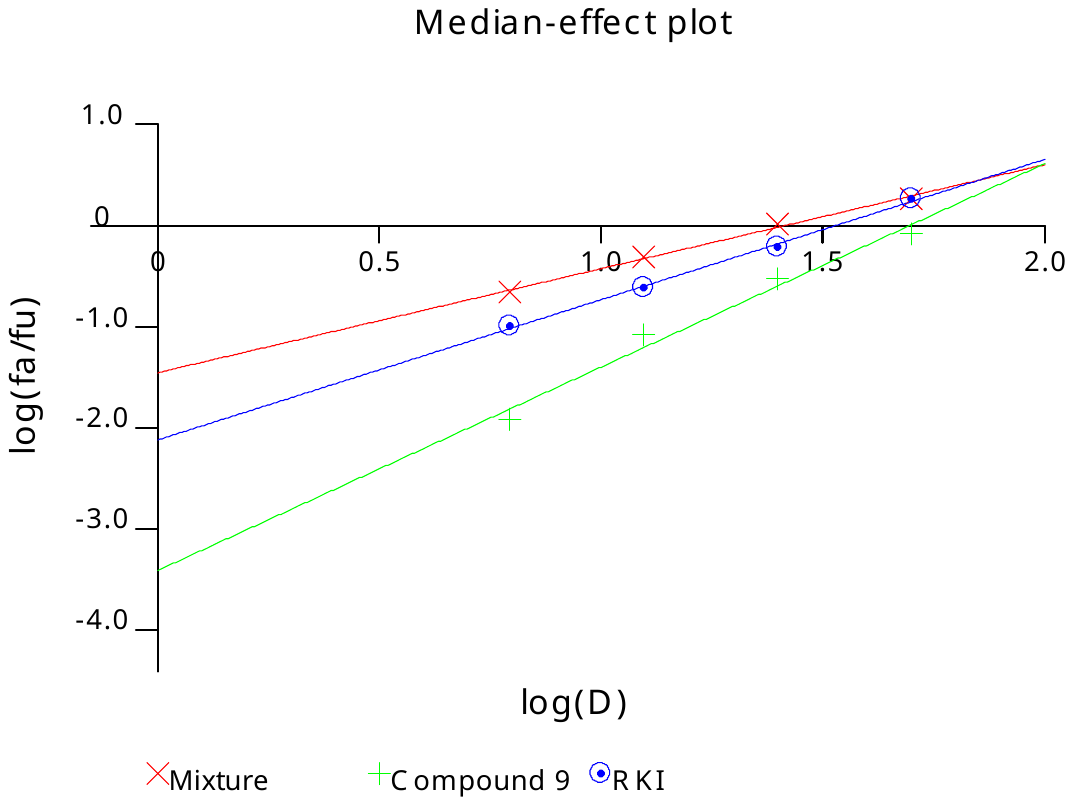

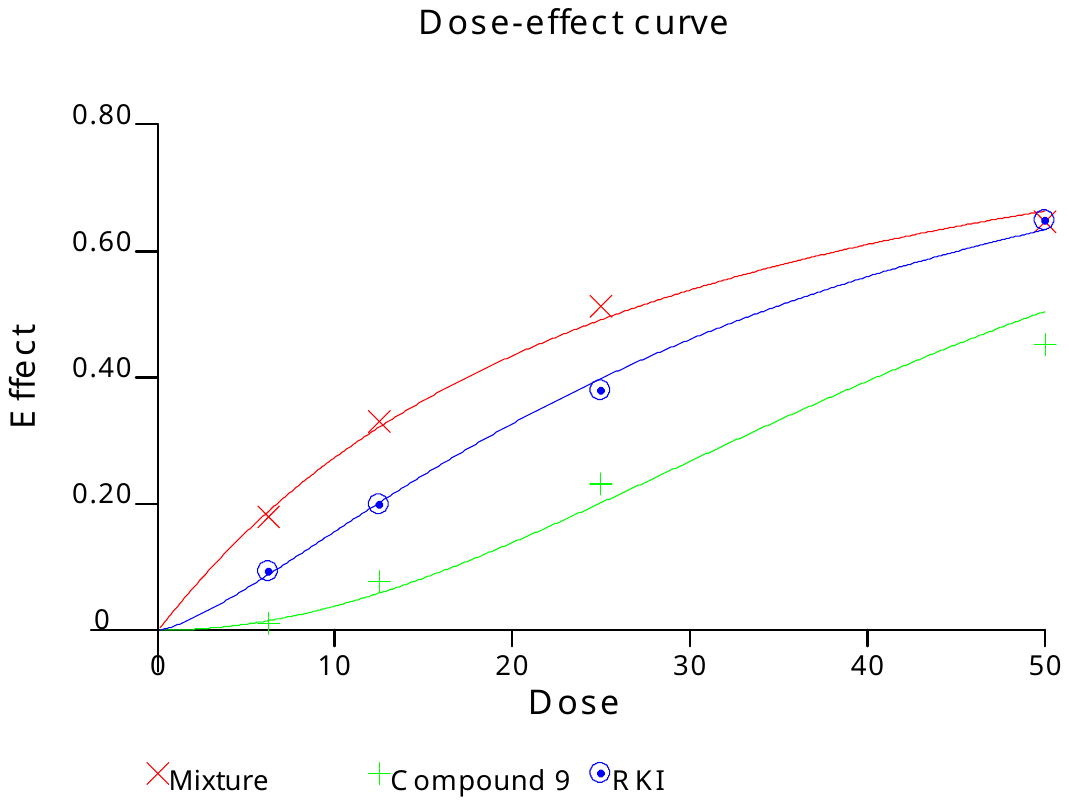

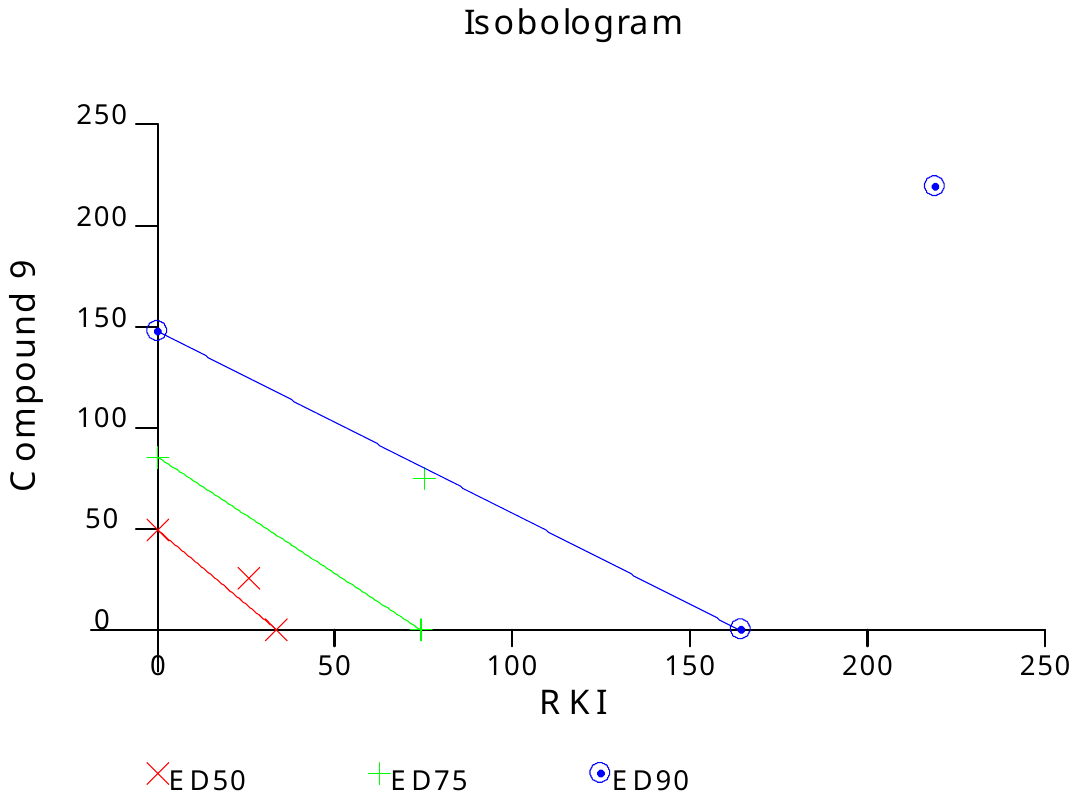

**MiaPaCa2 cell line**

|  | Drug concentration | | | |
| --- | --- | --- | --- | --- |
| Drug | I | II | III | IV |
| HDACi (**6b**) | 71.49±11.91 | 41.62±11.91 | 23.06±9.75 | 16.27±5.34 |
| Fingolimod | 79.98±10.36 | 48.31±12.04 | 20.57±4.85 | 20.33±5.37 |
| **6b** + Fingolimod | 89.28±4.05 | 60.81±20.19 | 34.11±14.58 | 26.96±7.59 |
| Combination Index (CI) | 0.780 | 1.422 | 1.648 | 1.070 |
| Interaction **6b** + Fingolimod | + + | − − | − − − | ± |

Compound 6 (µM): I=50; II=25; III=12.5; IV=6.25

Fingolimod (µM): I=8; II=4; III=2; IV=1

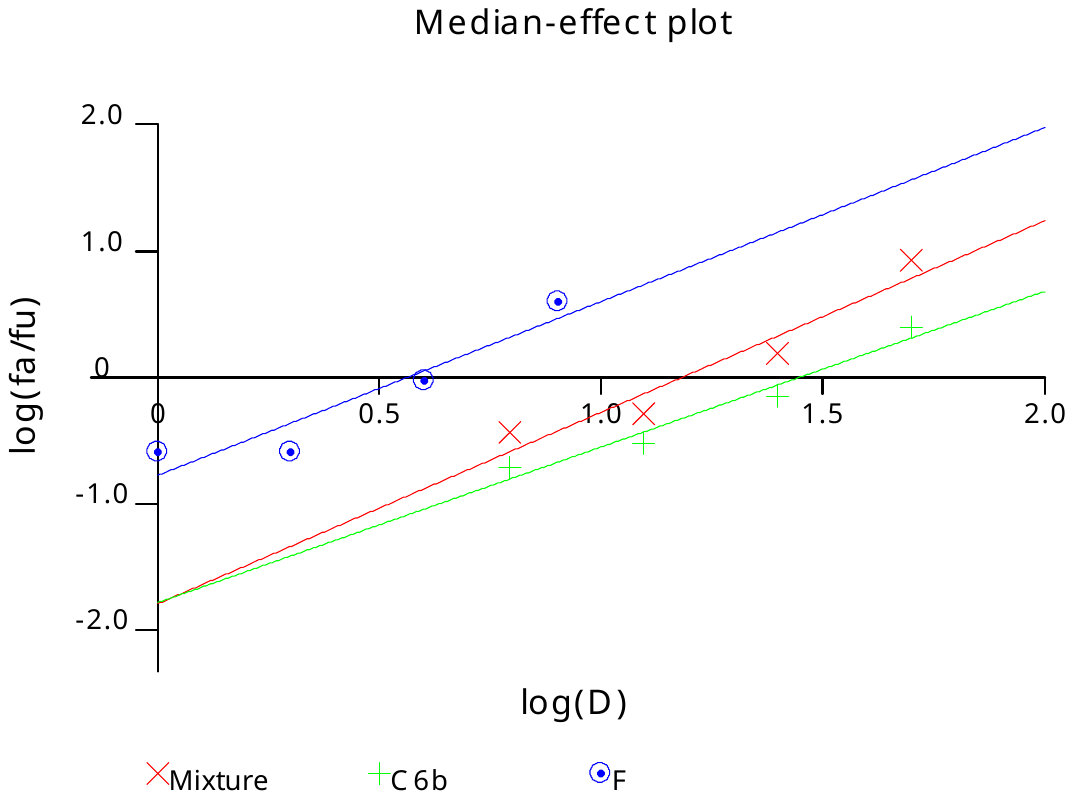

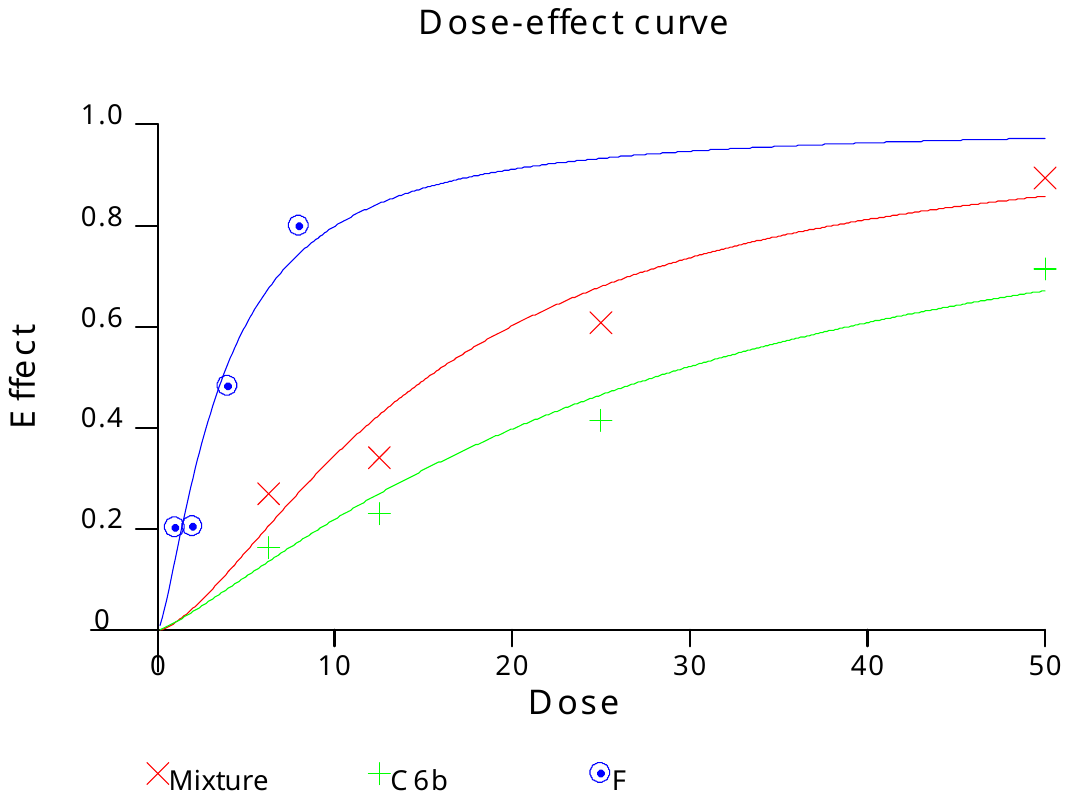

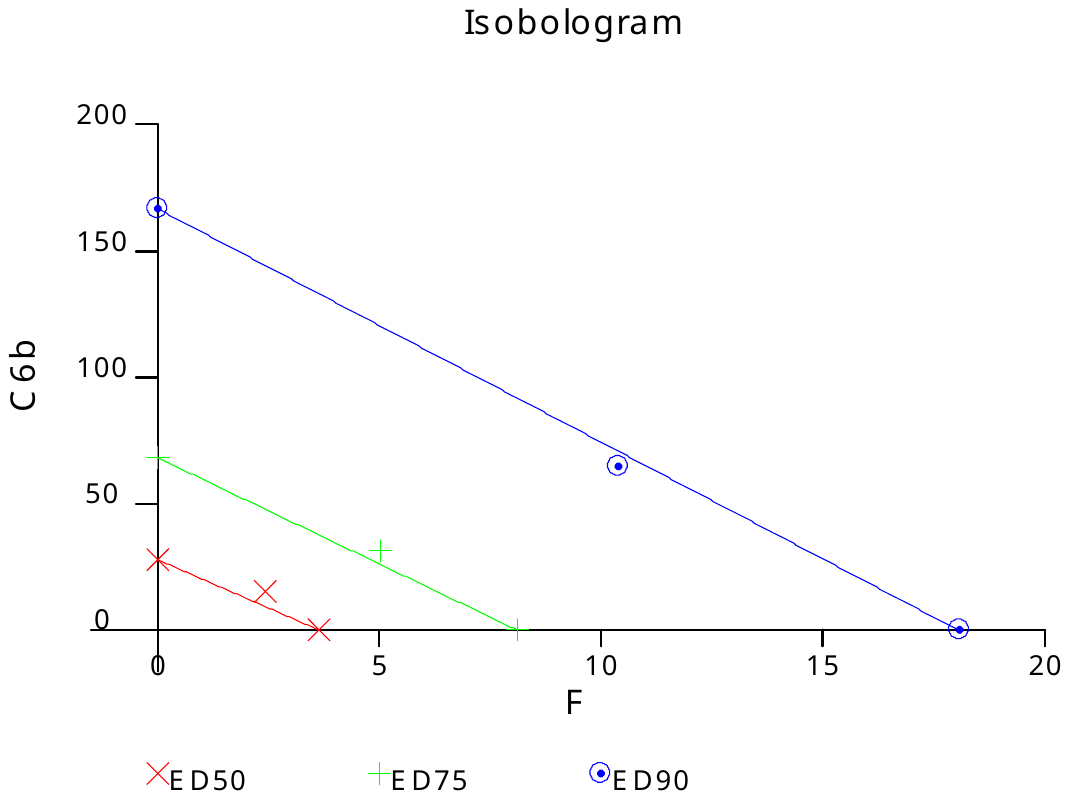

**MiaPaCa2 cell line**

|  | Drug concentration | | | |
| --- | --- | --- | --- | --- |
| Drug | I | II | III | IV |
| HDACi (**8b**) | 81.22±0.96 | 68.72±2.69 | 36.08±3.29 | 16.72±4.28 |
| Fingolimod | 87.22±2.01 | 52.02±2.47 | 23.73±3.92 | 22.83±1.89 |
| **8b** + Fingolimod | 91.31±10.08 | 65.06±1.92 | 38.98±5.26 | 26.55±2.94 |
| Combination Index (CI) | 1.188 | 1.836 | 1.852 | 1.338 |
| Interaction **8b** + Fingolimod | − − − | − − − | − − − | − − |

Compound 8 (µM): I=10; II=5; III=2.5; IV=1.25

Fingolimod (µM): I=8; II=4; III=2; IV=1

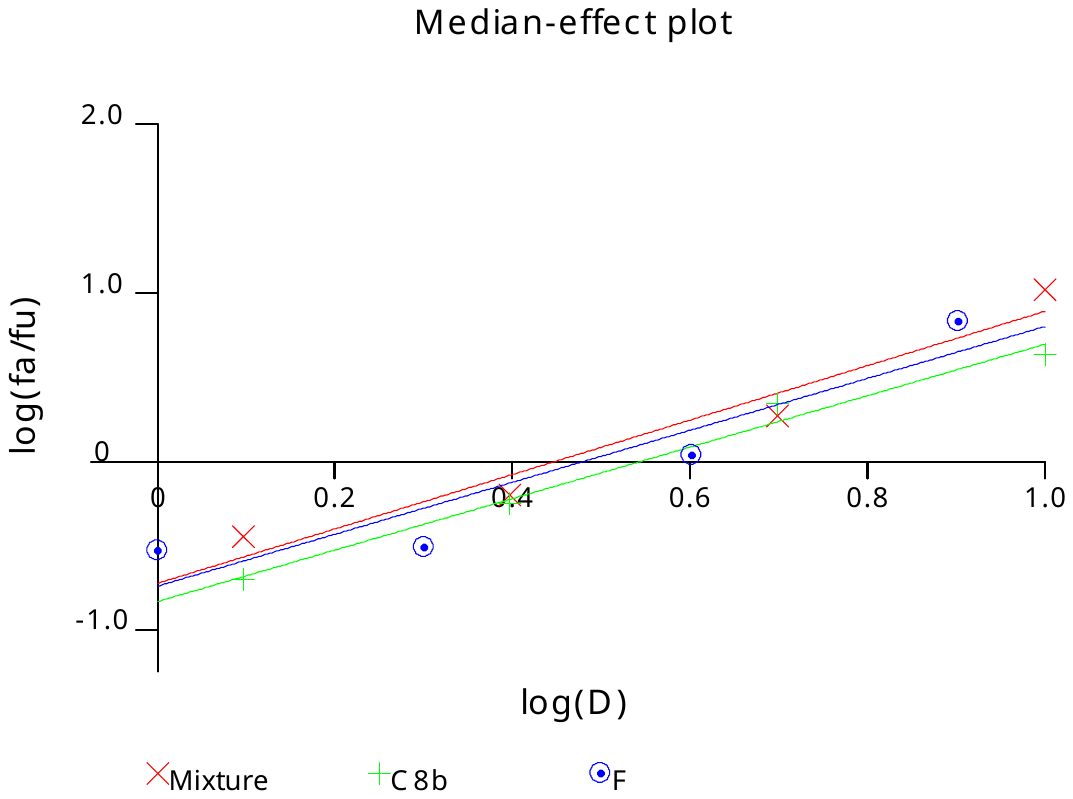

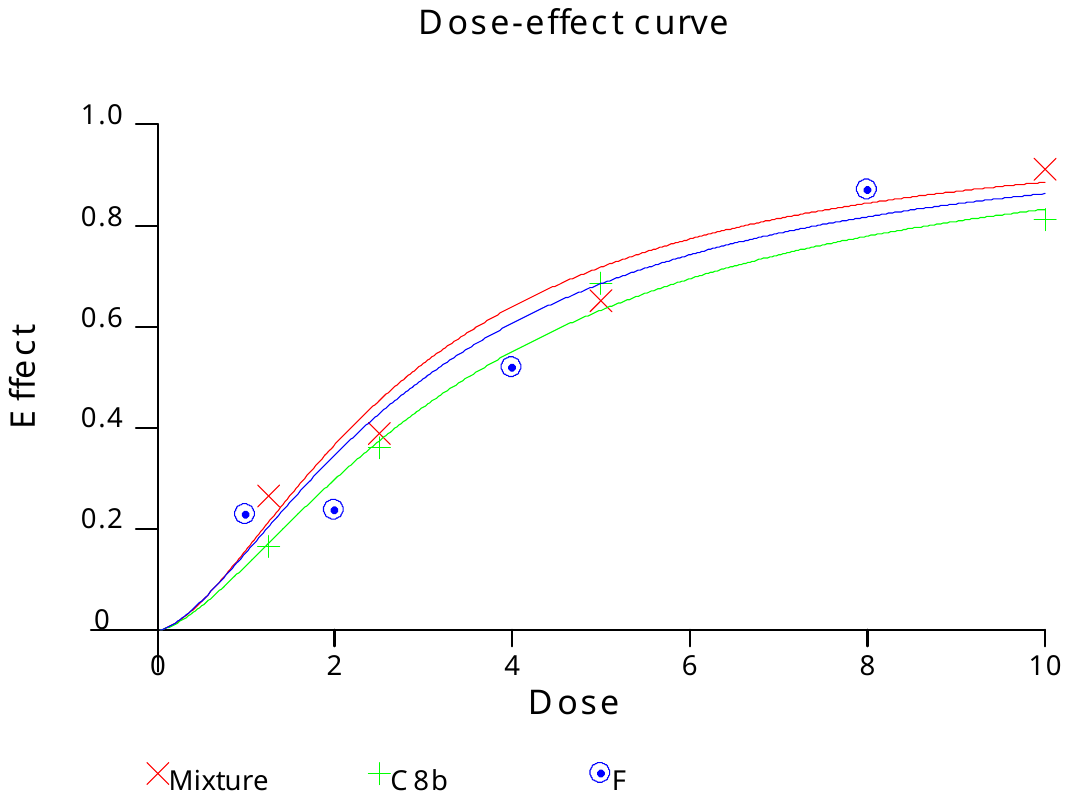

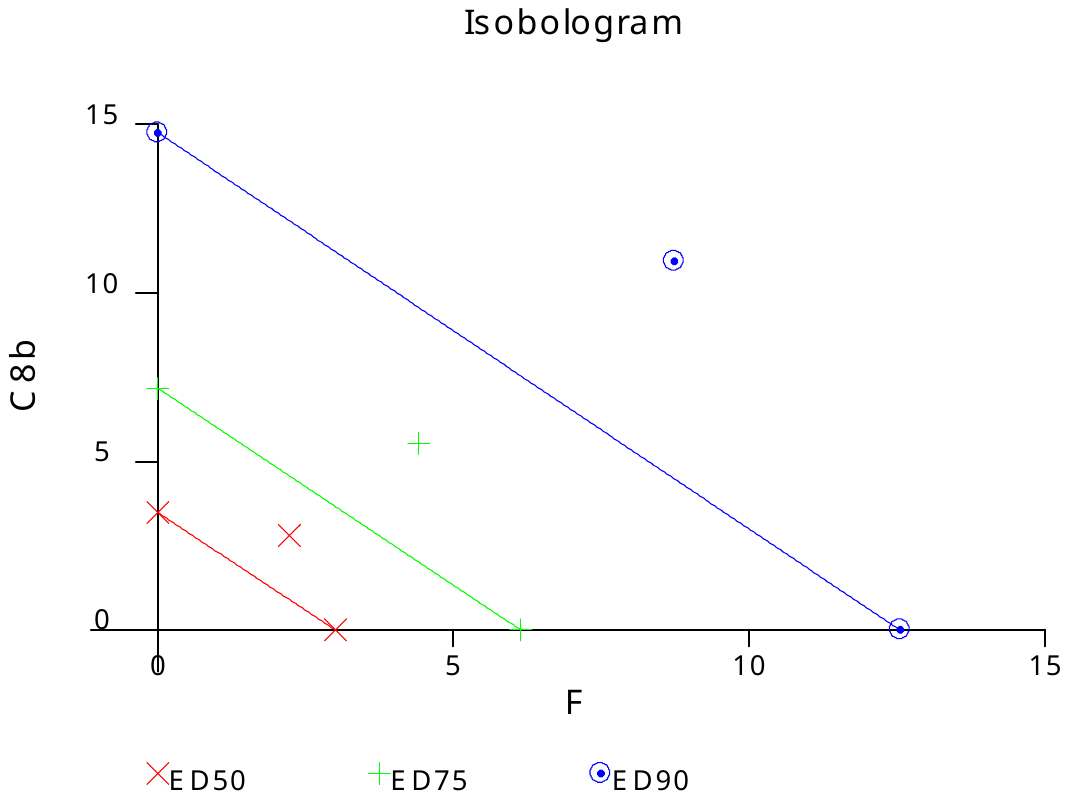

**MiaPaCa2 cell line**

|  | Drug concentration | | | |
| --- | --- | --- | --- | --- |
| Drug | I | II | III | IV |
| HDACi (**9b**) | 83.61±0.27 | 69.42±11.34 | 53.31±17.80 | 44.84±11.21 |
| Fingolimod | 76.58±4.48 | 63.82±6.81 | 43.78±12.65 | 35.34±6.99 |
| **9b** + Fingolimod | 89.11±1.57 | 73.59±12.40 | 55.57±18.16 | 47.22±9.51 |
| Combination Index (CI) | 0.870 | 1.449 | 1.773 | 1.293 |
| Interaction **9b** + Fingolimod | + | − − | − − − | − − |

Compound 9 (µM): I=50; II=25; III=12.5; IV=6.25

Fingolimod (µM): I=8; II=4; III=2; IV=1

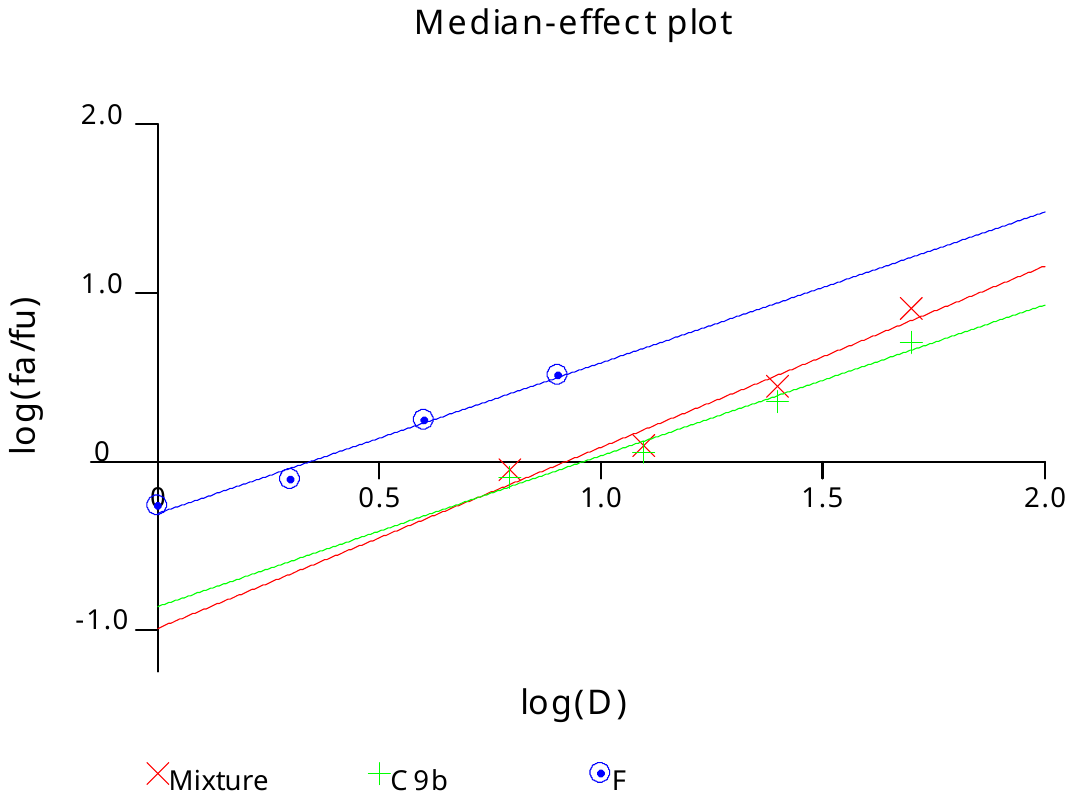

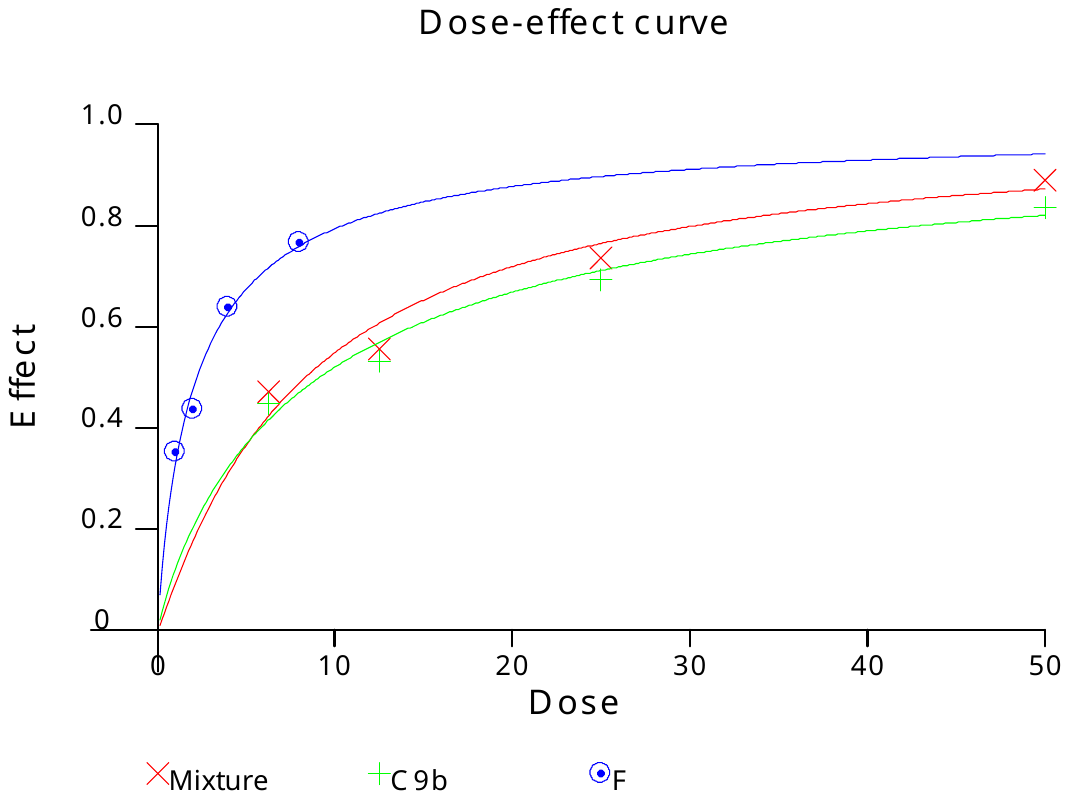

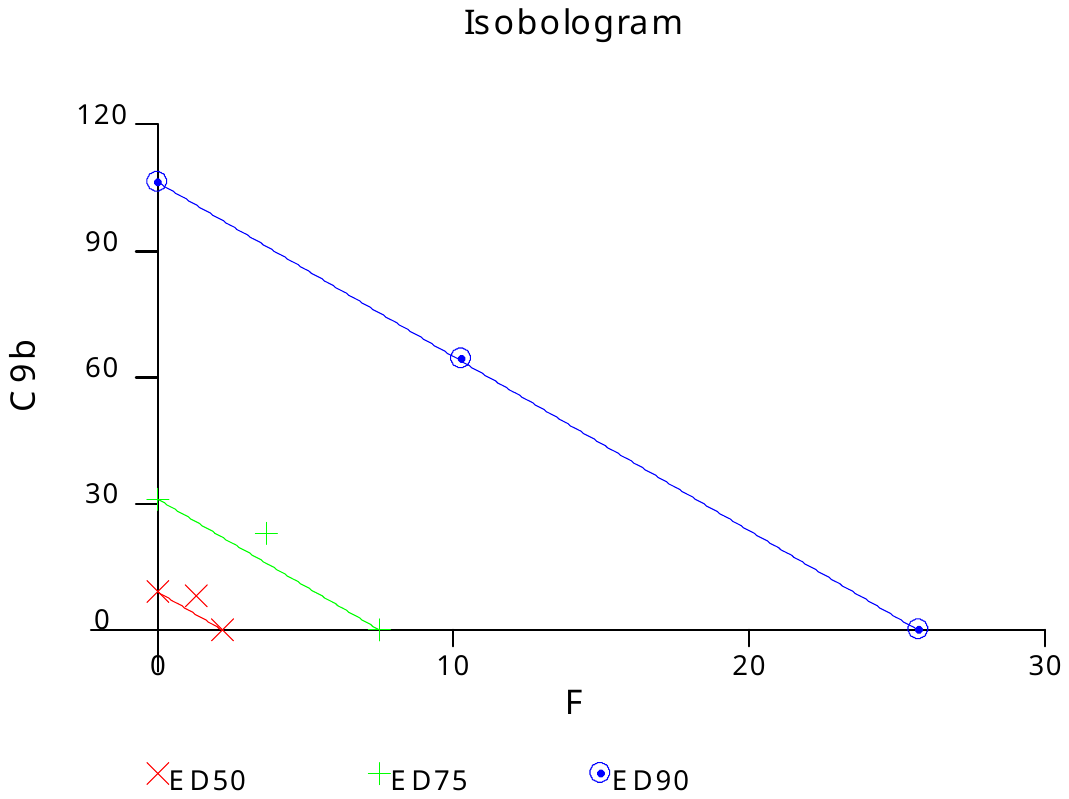

**Panc-1 cell line**

|  | Drug concentration | | | |
| --- | --- | --- | --- | --- |
| Drug | I | II | III | IV |
| HDACi (**6b**) | 63.68±4.70 | 25.29±6.71 | 23.93±6.23 | 10.30±1.96 |
| Fingolimod | 79.18±1.45 | 40.09±4.97 | 24.56±1.24 | 18.99±0.95 |
| **6b** + Fingolimod | 84.96±2.05 | 67.22±5.98 | 37.87±5.71 | 21.98±0.78 |
| Combination Index (CI) | 0.847 | 0.945 | 1.232 | 1.141 |
| Interaction **6b** + Fingolimod | + + | ± | − − | − |

Compound 6 (µM): I=100; II=50; III=25; IV=12.5

Fingolimod (µM): I=12; II=6; III=3; IV=1.5

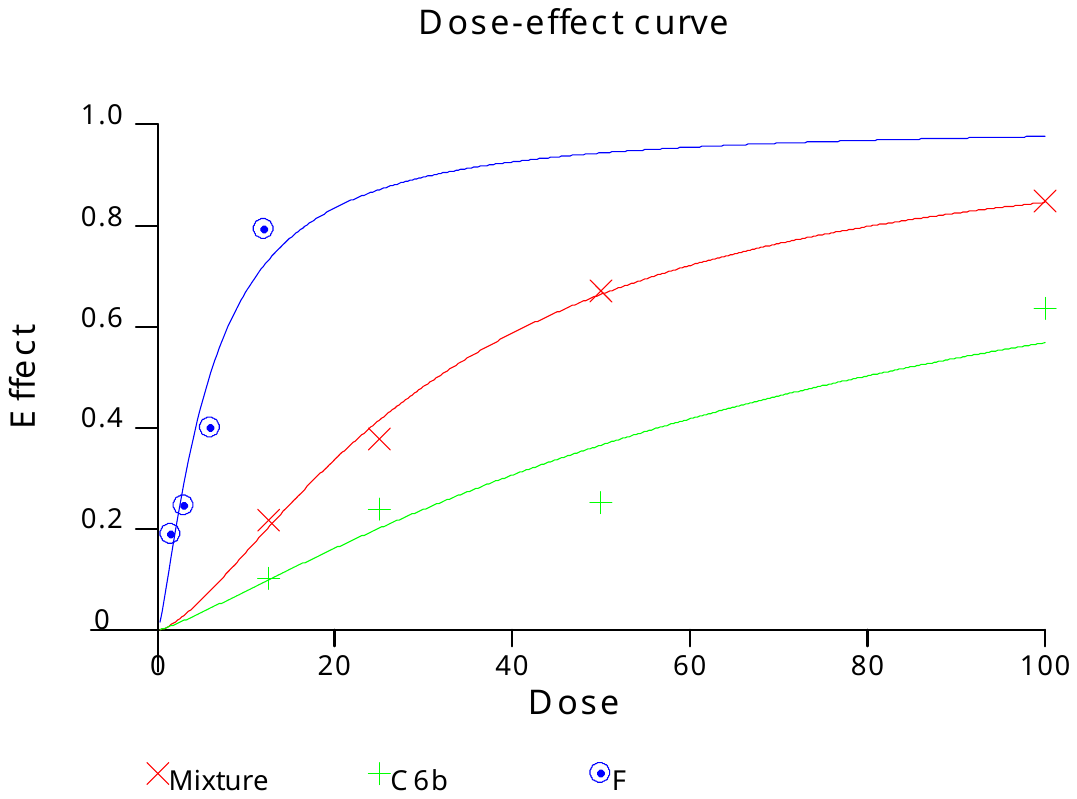

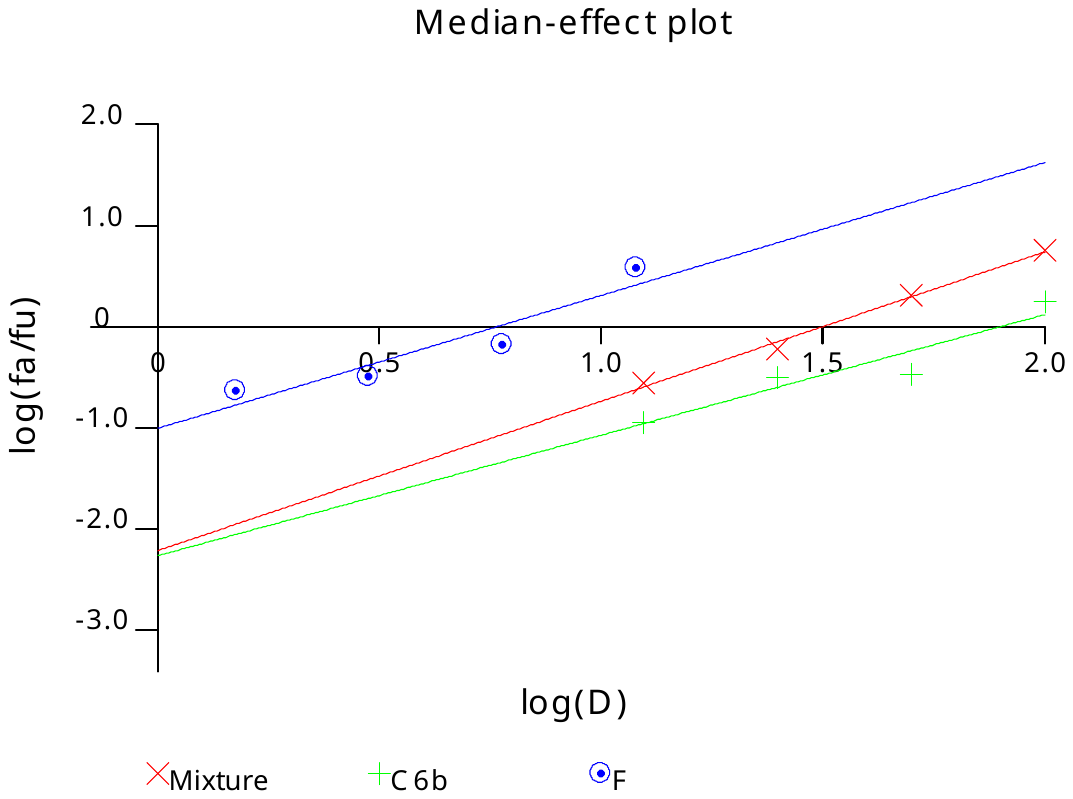

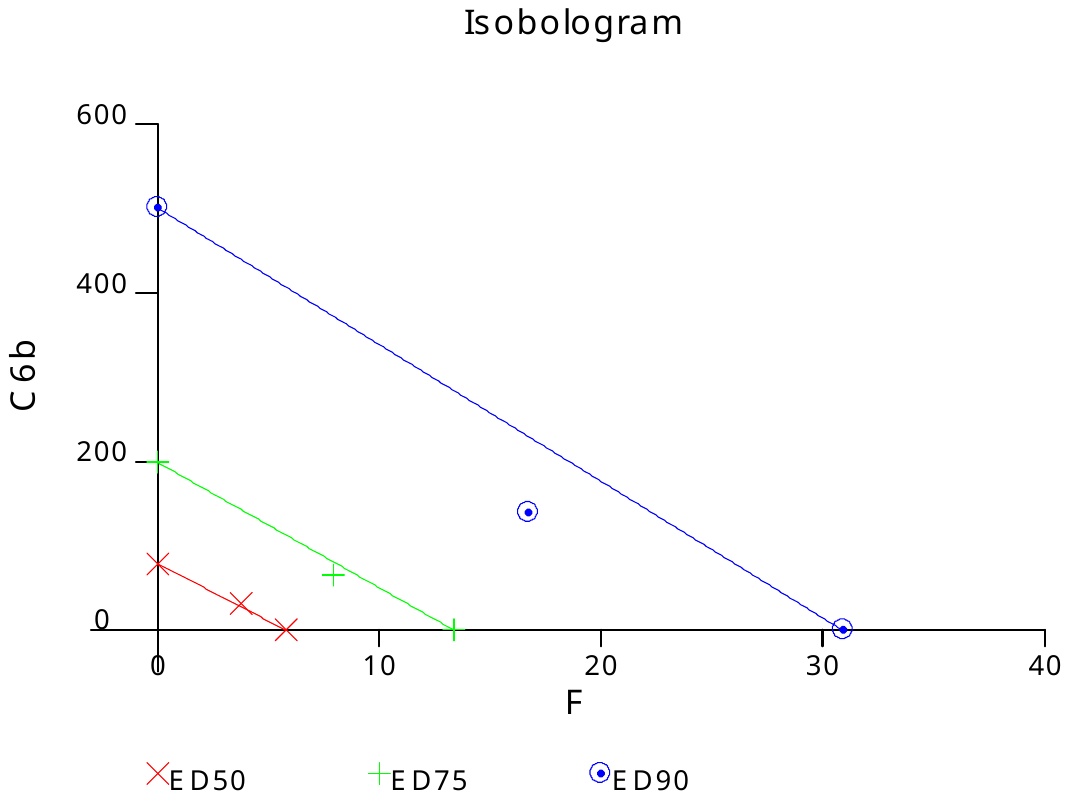

**Panc-1 cell line**

|  | Drug concentration | | | |
| --- | --- | --- | --- | --- |
| Drug | I | II | III | IV |
| HDACi (**8b**) | 58.10±0.24 | 23.86±9.24 | 4.77±2.69 | 4.68±2.33 |
| Fingolimod | 68.24±4.62 | 41.81±5.37 | 19.44±10.97 | 12.27±8.75 |
| **8b** + Fingolimod | 74.74±5.61 | 30.91±6.84 | 16.60±7.22 | 9.42±5.57 |
| Combination Index (CI) | 1.267 | 1.806 | 2.027 | 1.592 |
| Interaction **8b** + Fingolimod | − − | − − − | − − − − | − − − |

Compound 8 (µM): I=20; II=10; III=5.0; IV=2.5

Fingolimod (µM): I=12; II=6; III=3; IV=1.5

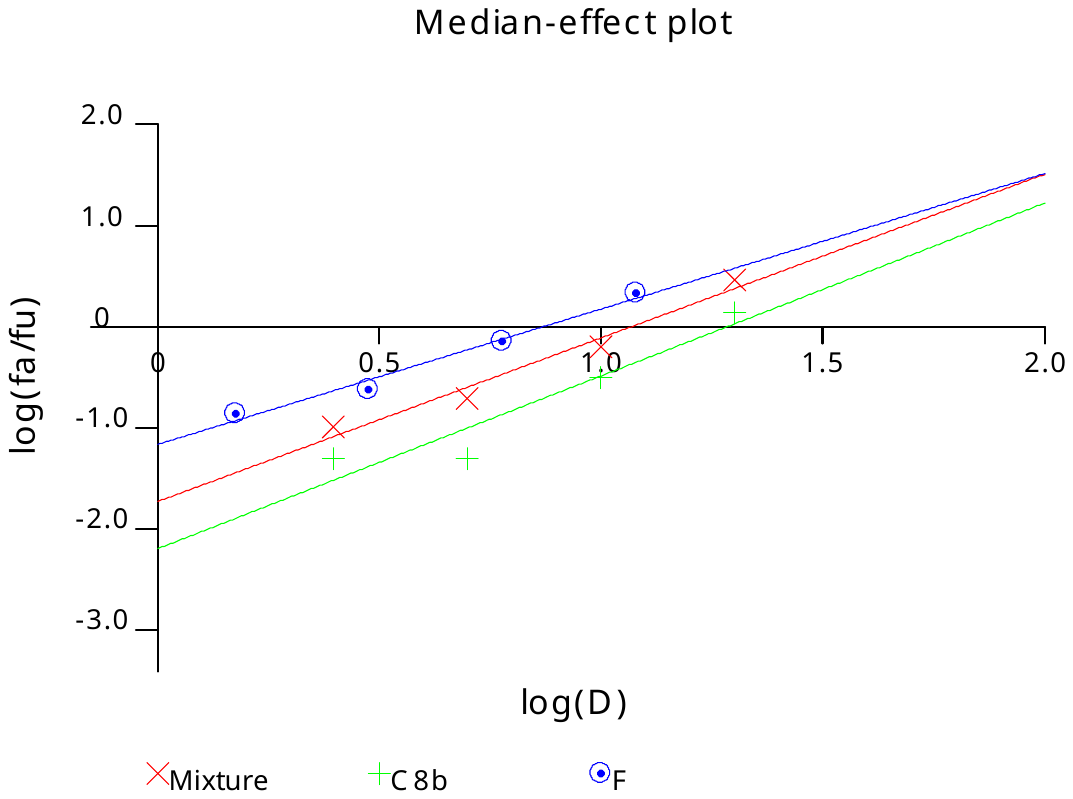

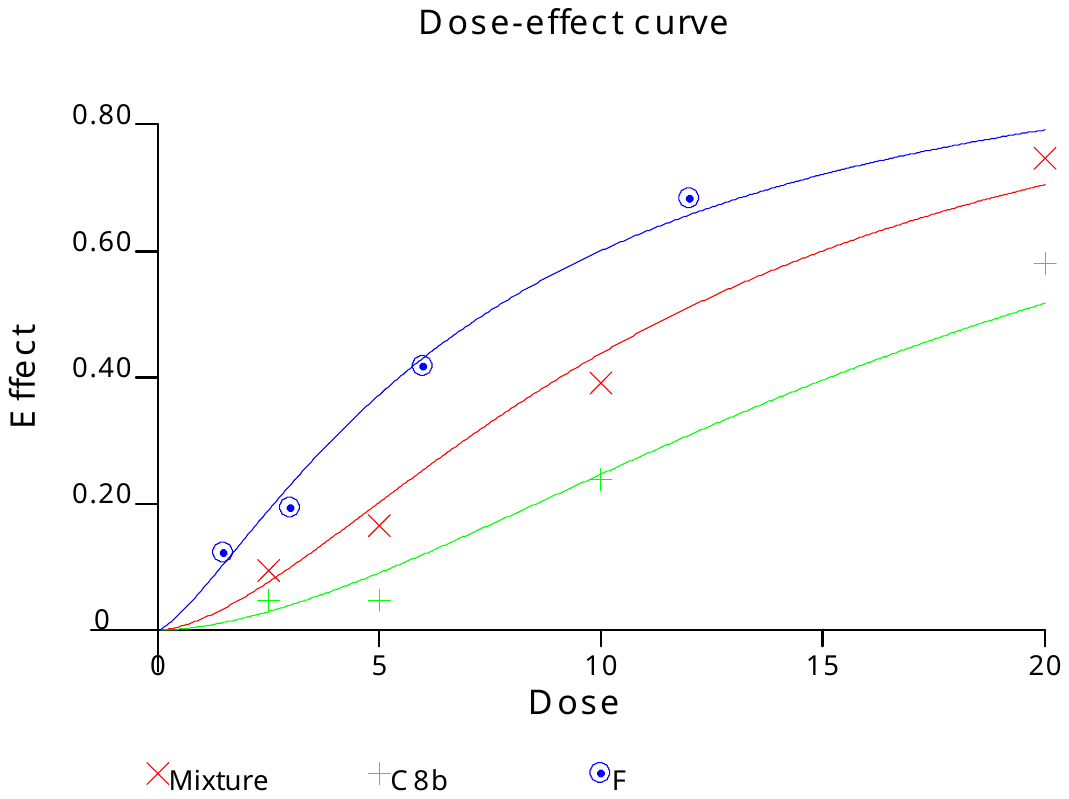

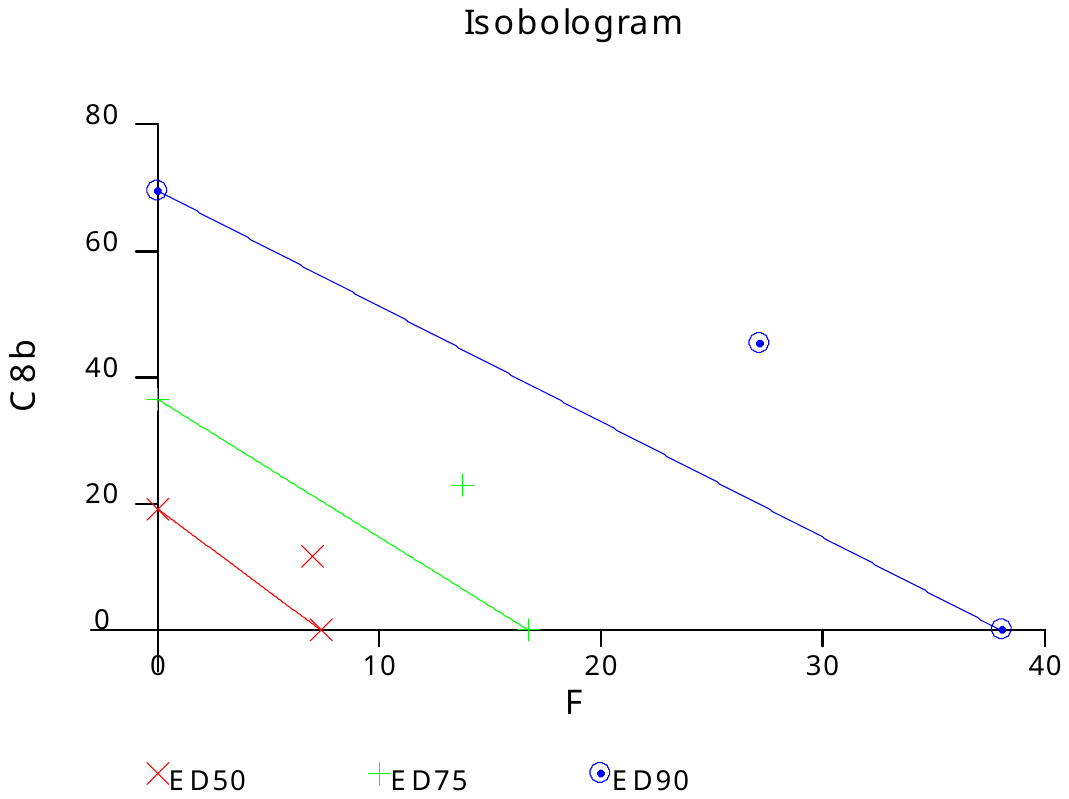

**Panc-1 cell line**

|  | Drug concentration | | | |
| --- | --- | --- | --- | --- |
| Drug | I | II | III | IV |
| HDACi (**9b**) | 61.28±6.91 | 50.93±3.83 | 40.71±3.74 | 24.03±5.73 |
| Fingolimod | 62.10±4.94 | 50.04±3.20 | 31.21±11.85 | 21.17±3.33 |
| **9b** + Fingolimod | 76.50±5.14 | 66.12±10.76 | 53.41±18.12 | 33.34±10.07 |
| Combination Index (CI) | 0.911 | 0.848 | 0.810 | 1.120 |
| Interaction **9b** + Fingolimod *^b^* | ± | + + | + + | − |

Compound 9 (µM): I=50; II=25; III=12.5; IV=6.25

Fingolimod (µM): I=12; II=6; III=3; IV=1.5

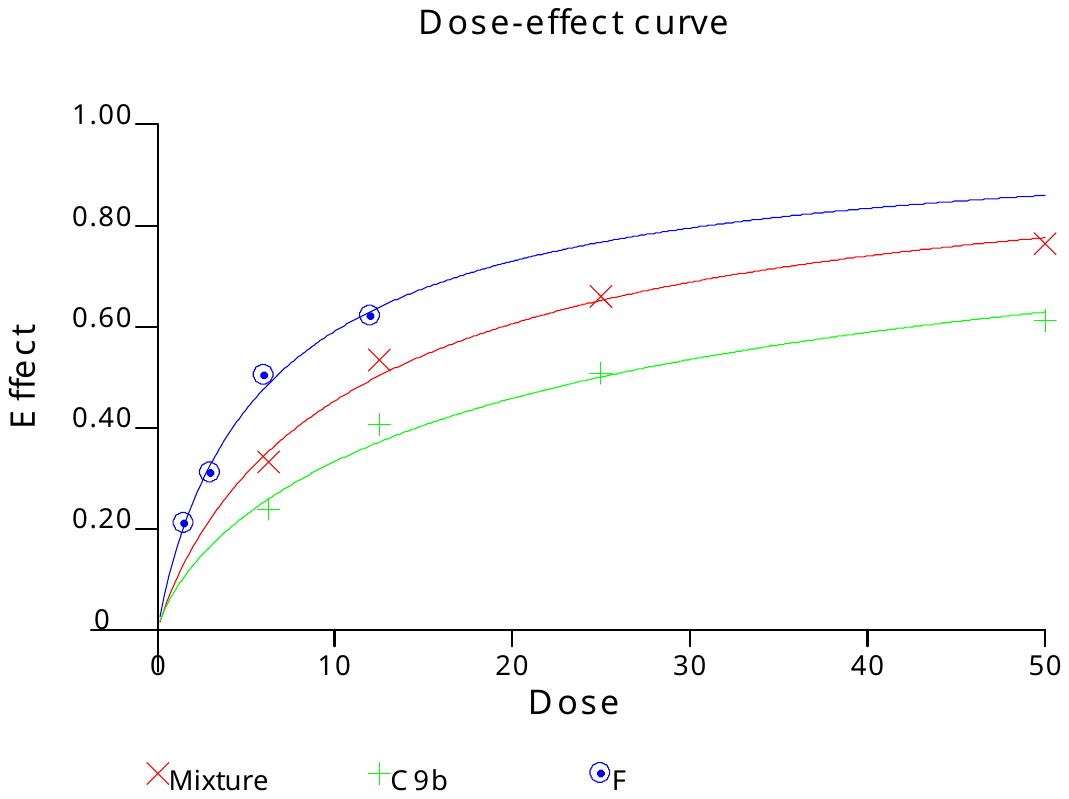

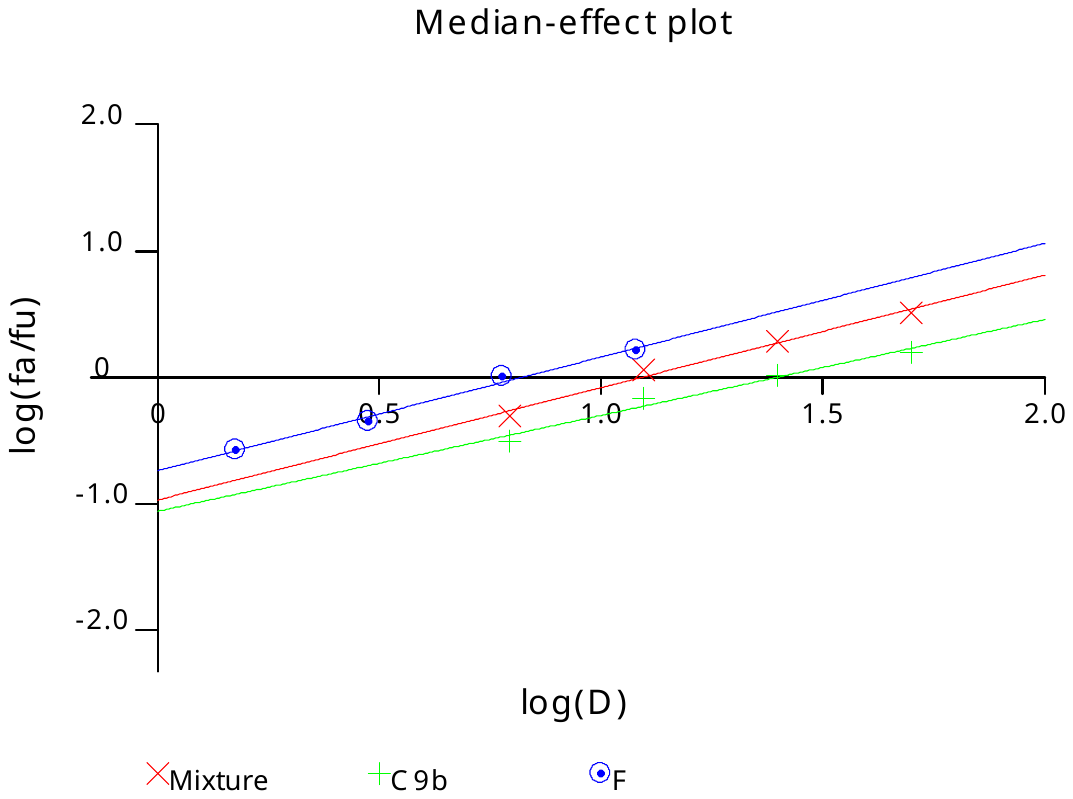

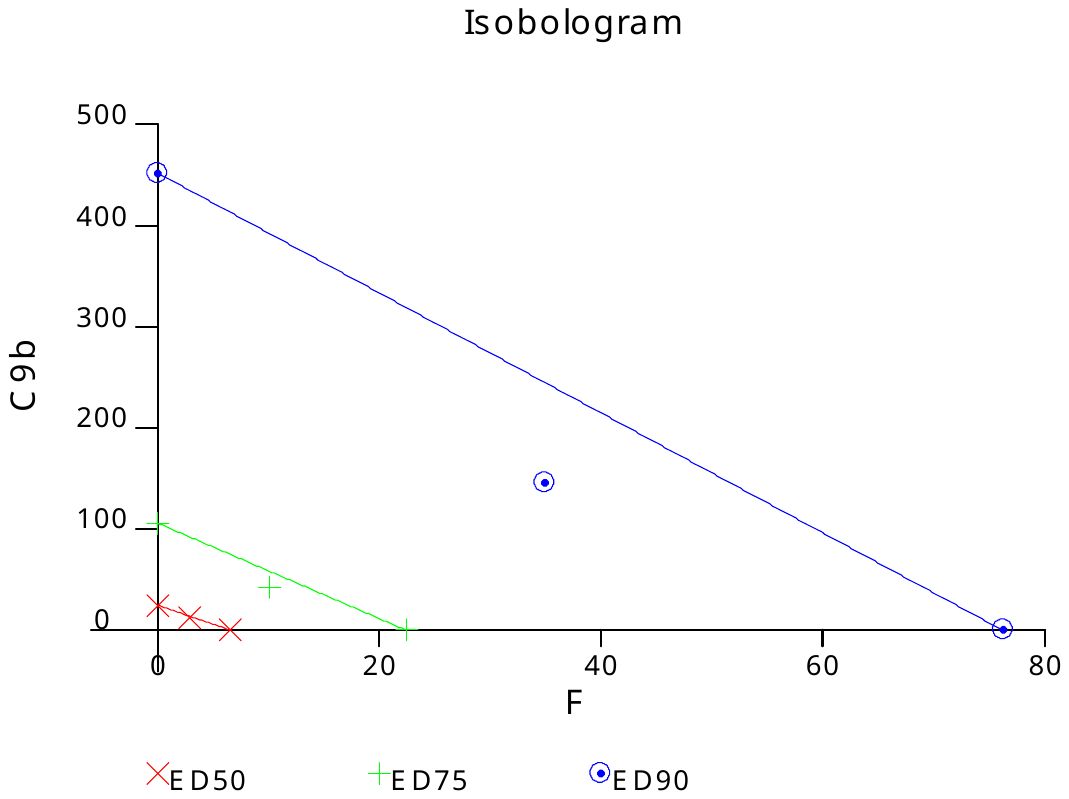

**Legend**

**Symbol	Description**

+ + + + +	Very strong synergism

**+ + +**	Synergism

**+ +**	Moderate synergism

+	Slight synergism

±	Nearly additive

**–**	Slight antagonism

**– –**	Moderate antagonism

**– – –**	Antagonism

– – – –	Strong antagonism

– – – – –	Very strong antagonism
